# Disparate roles for *C. elegans* DNA translocase paralogs RAD-54.L and RAD-54.B in meiotic prophase germ cells

**DOI:** 10.1101/2022.12.12.520157

**Authors:** Kei Yamaya, Bin Wang, Nadin Memar, Arome Solomon Odiba, Alexander Woglar, Anton Gartner, Anne M. Villeneuve

**Affiliations:** Department of Developmental Biology, Stanford University School of Medicine, Stanford, California, USA; State Key Laboratory of Non-food Biomass and Enzyme Technology, Guangxi Academy of Sciences, 530007 Nanning, China; IBS Center for Genomic Integrity and Department for Biological Sciences, Ulsan National Institute of Science and Technology, Ulsan, Korea; Swiss Institute for Experimental Cancer Research (ISREC), School of Life Sciences, Swiss Federal Institute of Technology Lausanne (EPFL), Lausanne, Switzerland (current affiliation); Department of Genetics, Stanford University School of Medicine, Stanford, California, USA

**Author notes:** Corresponding author: Anne M. Villeneuve.

## Abstract

RAD54 family DNA translocases partner with RAD51 recombinases to ensure stable genome inheritance, exhibiting biochemical activities both in promoting recombinase removal and in stabilizing recombinase association with DNA. Understanding how such disparate activities of RAD54 paralogs align with their biological roles is an ongoing challenge. Here we investigate the *in vivo* functions of *C. elegans* RAD54 paralogs RAD-54.L and RAD-54.B during meiotic prophase, revealing distinct contributions to the dynamics of RAD-51 association with DNA and to the progression of meiotic double-strand break repair (DSBR). While RAD-54.L is essential for RAD-51 removal from meiotic DSBR sites to enable recombination progression, RAD-54.B is largely dispensable for meiotic DSBR. However, RAD-54.B is required to prevent hyperaccumulation of RAD-51 on unbroken DNA during a key meiotic sub-stage when protein kinase CHK-2 is active. Moreover, DSB-independent hyperaccumulation of RAD-51 foci in the absence of RAD-54.B is RAD-54.L-dependent, revealing a hidden activity of RAD-54.L in promoting promiscuous RAD-51 association that is antagonized by RAD-54.B. We propose a model wherein a division of labor among RAD-54 paralogs allows germ cells to ramp up their capacity for efficient homologous recombination that is crucial to successful meiosis while counteracting potentially deleterious effects of unproductive RAD-51 association with unbroken DNA.

## Introduction

Homologous recombination plays a central role in in ensuring the faithful inheritance of genomes during meiosis, the specialized cell division program by which diploid organisms generate haploid gametes. In most sexually reproducing organisms, the formation of at least one crossover (CO) between each chromosome pair is crucial for faithful segregation of homologous chromosomes at the first meiotic division. To produce this obligate CO, many programmed double-strand DNA breaks (DSBs) are introduced into the genome by the meiosis-specific SPO11 protein complex and are subsequently repaired through homologous recombination. In most organisms studied, only a subset of DSB repair (DSBR) sites mature into COs; COs are often limited to one or a few per chromosome pair (or per chromosome arm), with the majority of introduced DSBs being repaired as non-COs (NCOs) (Gray and Cohen, 2016). Thus, both DSB formation and repair occur within a highly regulated framework that must simultaneously ensure both CO formation and restoration of genome integrity.

Recombinases play crucial roles in the early steps of homologous recombination. Following induction and processing of DSBs to yield 3’ single-stranded overhanging ends, recombinases are recruited to DSBR sites and form nucleoprotein filaments on the resected 3’ ssDNA, where they promote the search for a homologous DNA repair template and subsequently facilitate strand invasion into the template DNA duplex (Ceballos and Heyer, 2011; Gartner and Engebrecht, 2022). Most eukaryotes have two recombinases, RAD51, which is widely expressed, and DMC1, which is specialized to function in meiotic recombination, reflecting an ancient duplication of the ancestral recombinase gene early in the eukaryotic lineage (Bishop et al., 1992; Shinohara et al., 1992; Villeneuve and Hillers, 2001). However, DMC1 has been lost in several eukaryotic lineages, including many Hymenopteran and Dipteran insects and the nematode sublineage that includes the Caenorhabditis genus and its close relatives (Schurko et al., 2010; Villeneuve and Hillers, 2001). Thus, RAD-51 is the sole recombinase acting in both mitotically-cycling cells and meiotic prophase nuclei in the *C. elegans* germ line.

Eukaryotic recombinases RAD51 and DMC1 function in partnership with RAD54 family DNA translocases, which are members of the Snf2/Swi2 group of ATP-dependent DNA motors (Ceballos and Heyer, 2011; Crickard, 2021). RAD54 family proteins have a conserved C-terminal ATPase/helicase-like domain that promotes translocation along double-stranded (ds) DNA and a more variable N-terminal domain that mediates interactions with recombinase partners and is required for multiple aspects of RAD54 function (Ceballos and Heyer, 2011; Crickard, 2021).

RAD54L orthologs, represented by Rad54 in *S. cerevisiae* and previously called RAD54 or RAD54A in the mammalian literature, have been studied extensively through both biochemical and cell-based assays (Ceballos and Heyer, 2011; Crickard, 2021). *In vitro*, RAD54(L) has been demonstrated to have dsDNA-stimulated ATPase activity, ATP-dependent dsDNA translocase activity, and ATP-dependent chromatin remodeling activity. Further, numerous *in vitro* assays have demonstrated several activities relevant to RAD51 recombinase-mediated homologous recombination. These include stabilization of RAD51 association with ssDNA, removal of RAD51 filaments from dsDNA (mediated by RAD54 translocation), and stimulation of RAD51-mediated formation of D-loops (key DSBR intermediates resulting from successful strand invasion), with this latter activity likely reflecting contributions of both RAD51 stabilization and removal activities (Ceballos and Heyer, 2011; Crickard, 2021). In addition, *in vitro,* single-molecule imaging assays suggest that the RAD51-filament stabilizing and dsDNA translocase activities of RAD54(L) may act together to enhance the efficiency of RAD51-mediated homology search by enabling motor-driven one-dimensional translocation of RAD51-ssDNA nucleoprotein filaments along dsDNA (Crickard et al., 2020b; Meir et al., 2022). *In vivo* studies support the biological importance of a close functional partnership between RAD54L orthologs and RAD51 in maintaining genome integrity. *S. cerevisiae* Rad54 is required for DNA repair after damage in mitotic cells, with mutants exhibiting a phenotype identical to *rad51* mutants, and mammalian RAD54 is required for homologous recombination in mammalian cells (Ceballos and Heyer, 2011). Further, cytological evidence from mammalian cells supports RAD54 having both an ATP-independent activity that promotes efficient recruitment of RAD51 to DSBR sites and an ATP-dependent activity that promotes RAD51 removal (Agarwal et al., 2011; Tan et al., 1999). In *C. elegans* meiocytes, RAD-54.L (formerly known as RAD-54) has been shown to be essential for removal of RAD-51 from DSBR sites (Mets and Meyer, 2009; Nadarajan et al., 2021; Roelens et al., 2019b; Saito et al., 2012).

Many eukaryotes have a second RAD54 paralog, exemplified by Rdh54/Tid1 in *S. cerevisiae* and RAD54B in mammals (Crickard, 2021). The phylogenetic distribution of RAD54 paralogs is consistent with the possibility that separate RAD54L/Rad54 and RAD54B/Rdh54 paralogs originated from an ancient gene duplication that predated the divergence of plants, animals, and fungi (Crickard, 2021; Schurko et al., 2010;

WormBase, WS286). In contrast to RAD54L, where clear orthologs are present through the eukaryotic lineage, apparent RAD54B/Rdh54 orthologs are absent from many lineages and exhibit greater sequence diversification than RAD54L in lineages where they are present. We speculate that duplication of an ancestral RAD54 gene to yield separate RAD54L and RAD54B paralogs may have been coupled to duplication of the ancestral recombinase RAD51 gene to yield the meiosis-specific recombinase DMC1. However, this is neither an essential nor exclusive partnership, as RAD54B is absent from many plant lineages and some arthropod lineages that retain DMC1 (Schurko et al., 2010), and RAD54B is present in Caenorhabditis nematodes despite the loss of DMC1 (WormBase, WS286).

Genetic and biochemical studies have provided evidence for both distinct and partially overlapping roles for RAD54L and RAD54B homologs. Biochemically, Rad54 and Rdh54 have been shown to be very similar, with some differences in their ATPase activity and translocation velocity and processivity (Ceballos and Heyer, 2011; Crickard, 2021). However, genetic evidence indicates a significant division of labor between the two paralogs: *e.g.* the *S. cerevisiae rad54* mutant primarily exhibits defects in mitotic DSBR, while the *rdh54* mutant primarily exhibits defects in meiotic DSBR (Shinohara et al., 1997). Consistent with these divergent mutant phenotypes, biochemical evidence supports Dmc1 preferentially functioning together with Rdh54, and Rad51 preferentially functioning with Rad54 (Nimonkar et al., 2012). Despite this clear evidence for specialization, however, multiple studies also indicate partial functional redundancy between the two paralogs. For example, the yeast *rad54 rdh54* double mutant exhibits stronger defects in both mitosis and meiosis compared to either single mutant (Shinohara et al., 1997). Moreover, additional functions for *S. cerevisiae* Rdh54 and mammalian RAD54B have also been demonstrated in mitotically dividing cells where DMC1 is absent and RAD51 is the only recombinase. For example, yeast Rdh54 has been implicated in limiting the size of Rad51-mediated D-loop intermediates *in vivo* (Keymakh et al., 2022; Shah et al., 2020), and biochemical evidence indicates that the two RAD54 paralogs may occupy different sites on assembled presynaptic filaments (Crickard et al., 2020a), suggesting that both RAD54 paralogs may be deployed in different ways at the same DSBR site. Further, loss of either RAD54 paralog causes increased sensitivity to DNA damaging agents both in mouse embryonic stem cells and mice, with loss of both paralogs causing stronger sensitivity (Wesoly et al., 2006). Additionally, RAD54B orthologs have been shown to antagonize potentially toxic association of RAD51 with unbroken DNA in both mammalian cells and vegetatively growing yeast cells (Mason et al., 2015; Shah et al., 2010), paralleling a role for Rdh54 in antagonizing association of Dmc1 with unbroken DNA during yeast meiosis (Holzen et al., 2006). Thus, while multiple disparate biochemical activities have been identified for RAD54 family proteins, much remains to be learned regarding which activities are employed *in vivo* and how each paralog may contribute differentially to DSBR in different biological contexts.

Here, we investigate the *in vivo* biological roles of RAD54 family paralogs RAD-54.L and RAD-54.B during meiosis in the nematode *C. elegans*. We demonstrate that RAD-54.L and RAD-54.B make distinct contributions to meiotic homologous recombination and provide evidence for disparate activities in regulating RAD-51 recombinase. While RAD-54.L is essential for the progression of meiotic DSBR, RAD-54.B is largely dispensable for the completion of meiotic recombination and instead functions in inhibiting the promiscuous accumulation of RAD-51 on unbroken DNA. Unexpectedly, we found that hyperaccumulation of RAD-51 at unbroken DNA in the absence of RAD-54.B is dependent on RAD-54.L, a surprising *in vivo* demonstration of its activity to stabilize/promote RAD-51 binding, indicating that RAD-54.L can both promote and antagonize RAD-51 binding in the same cells. Taken together, our data suggest a model in which RAD-54.L is hyperactivated during meiotic prophase to accommodate an increased burden on the DSBR machinery imposed by programmed DSB induction. We propose that RAD-54.B counteracts the potentially deleterious effects of RAD-54.L hyperactivation by inhibiting ectopic RAD-51 binding to unbroken DNA.

## Results

### *rad-54.B* mutants undergo mostly normal meiosis but exhibit hyperaccumulation of RAD-51 foci

Our investigation of the roles of RAD-54.B was initiated by our identification of missense mutation *rad-54.B(gt3308)* in a genetic screen for *C. elegans* mutants exhibiting an increased sensitivity to ionizing radiation at the L1 larval stage (**Fig 1A**) (González-Huici et al., 2017). Given that the *C. elegans* RAD-54.B paralog RAD-54.L (previously known as RAD-54) is essential for DSBR during meiotic recombination, and that the *S. cerevisiae* RAD-54.B ortholog Rdh54/Tid1 contributes to meiotic recombination in budding yeast, we investigated potential effects of loss of *rad-54.B* function on progression and success of meiotic recombination.

**Fig 1.**
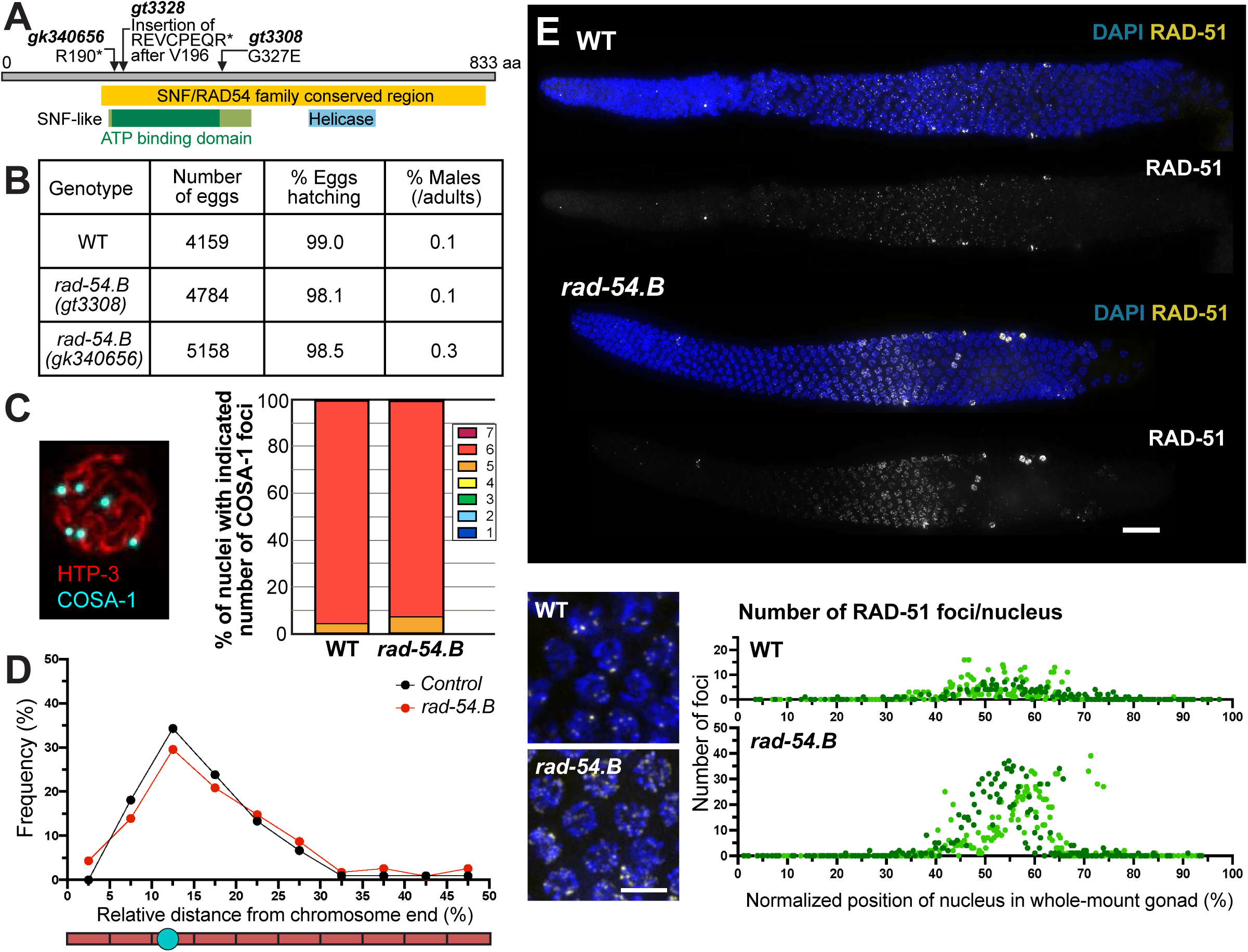
Loss of *rad-54.B* function is compatible with successful meiosis but causes temporary hyperaccumulation of RAD-51. A) Diagram of RAD-54.B protein with alterations caused by mutant alleles. Yellow box indicates region conserved across SNF/RAD54 family members. Green box indicates SNF2-like N-terminal domain, with dark green indicating the ATP-binding domain of helicase superfamily 1/2; blue box indicates C terminal helicase domain. Unless otherwise noted, experiments were conducted using the *gk340656* presumed null allele. B) Quantification of progeny viability and fraction of males among adult progeny, indicating successful meiotic chromosome segregation in *rad-54.B* mutants. Number of broods assayed: WT, n=13; *rad-54.B(gt3308)*, n=16; *rad-54.B(gk340656)*, n=16. C) Left: Example *meIs8[pie-1p::GFP::cosa-1::unc-119(+)]* nucleus immunostained for HTP-3 and GFP. Right: Quantification of COSA-1 foci in late pachytene nuclei from WT and *rad-54.B* gonads. Number of nuclei assayed: WT, n=251; *rad-54.B*, n=283. D) Distribution of positions of CO site foci of control (*meIs8; mnT12*) and *rad-54.B* (*meIs8; rad-54.B mnT12*) late pachytene nuclei on non-fusion chromosomes, represented as relative distance from the nearest chromosome end. Control, n=88 chromosomes; *rad-54.B*, n=100 chromosomes. E) Top: Max-projected images of WT and *rad-54.B* whole-mount gonads immunostained for RAD-51, showing elevated levels of RAD-51 foci detected in the *rad-54.B* mutant. Scale bar represents 20 μm. Bottom, left: Zoomed-in fields of nuclei in zone of peak RAD-51 accumulation in both WT and *rad-54.B* gonads. Bottom, right: Quantification of RAD-51 foci in WT and *rad-54.B* gonads. Each data point represents an individual nucleus, with relative position calculated as the percentage of the length between the distal tip (0%) and the end of pachytene zone (100%); light and dark colors represent the two different gonads used for quantification.

Several lines of evidence indicate that RAD-54.B is largely dispensable for successful meiosis in *C. elegans*. First, worms homozygous either for *rad-54.B(gt3308)* or for putative null allele *rad-54.B(gk340656)* exhibit nearly wild type (WT)-like levels of embryonic viability and male progeny (**Fig 1B**); this contrasts with expectations for *C. elegans* mutants defective in meiotic recombination, which exhibit a high rate of embryonic lethality (reflecting failure of DSBR or missegregation of autosomes) and/or a “high incidence of male” progeny (the Him phenotype, reflecting sex-chromosome mis-segregation). We also quantified COSA-1 foci, which mark crossover (CO) sites at the late pachytene stage of meiotic prophase. Six COSA-1 foci are typically observed in WT nuclei, corresponding to a single CO site for every chromosome pair (Yokoo et al., 2012). Six COSA-1 foci per nucleus were similarly detected in *rad-54.B(gk340656)* late-pachytene meiocytes, suggesting that COs are specified in normal numbers in this mutant (**Fig 1C**). Further, distributions of the positions of COSA-1 foci along chromosome axes were similar in control and mutant meiocytes (**Fig 1D**), and fusion-chromosome assays for CO patterning likewise did not detect differences between mutant and control (**Fig S2**). Together, these data suggest that the mechanisms required to initiate meiotic recombination and to promote and regulate the designation of meiotic CO sites are largely operational in *rad-54B* mutants.

Despite the above evidence for substantially successful CO formation and meiotic chromosome segregation in *rad-54.*B mutants, immunostaining for recombinase RAD-51 revealed a striking temporary hyperaccumulation of RAD-51 foci in two independently-derived *rad-54.*B presumed null mutants, *rad-54.B(gk340656)* and *rad-54.B(gt3328)* (**Fig 1E, S1**). In a WT whole-mount gonad, RAD-51 can be visualized as chromosome-associated foci in prophase nuclei starting at the transition zone and ending midway through pachytene (**Fig 1E, S1**). In contrast, both *rad-54.B* mutants exhibit hyperaccumulation of RAD-51 foci within a specific region of the germ line, starting at meiotic prophase onset and declining abruptly midway through prophase I (**Fig 1E, S1**). In subsequent experiments in this study, we use *rad-54.B(gk340656)*, a presumed null allele, unless otherwise specified.

We note that the temporary hyperaccumulation of RAD-51 observed in *rad-54.B* mutants contrast sharply with RAD-51 hyperaccumulation observed in *rad-54.L* mutants, in which RAD-51 foci continue to accumulate throughout meiotic progression (**Fig S1**). Given this difference in RAD-51 hyperaccumulation, we further investigated the progression of meiotic DSBR in these mutants.

### Meiotic DSBR progression is severely impaired in *rad-54.L* mutant but appears substantially normal in *rad-54.B* mutant

We investigated progression of meiotic DSBR in *rad-54.B* and *rad-54.L* mutants by simultaneous immunostaining for RAD-51 and MSH-5 in spread preparations that improve detection of chromosome-associated recombination proteins. MSH-5, which partners with MSH-4 to comprise the MutSg complex, is a meiosis-specific DSBR factor that initially localizes to numerous interhomolog recombination intermediates in early pachytene nuclei and becomes concentrated specifically at CO-designated sites upon transition to late pachytene (Woglar and Villeneuve, 2018; Yokoo et al., 2012). Time course analysis indicates that early pachytene MSH-5 foci in WT meiosis represent post-strand-exchange intermediates (Woglar and Villeneuve, 2018).

Our cytological analysis revealed several notable differences between the *rad-54.L* and *rad-54.B* mutants. First, abundant MSH-5 foci accumulate in *rad-54.B* early pachytene nuclei, albeit with an apparent delay (**Fig 2A**, **2B**). In contrast, numbers of MSH-5 foci in the *rad-54.L* mutant are greatly reduced relative to either WT or *rad-54.B* (**Fig 2A**, **2B**). Second, while MSH-5 foci and RAD-51 foci rarely colocalize in either WT or *rad-54.B* early pachytene nuclei despite increased numbers of RAD-51 foci in the *rad-54.B* mutant, the few MSH-5 foci observed in *rad-54.L* early pachytene nuclei frequently colocalize with RAD-51 (**Fig 2A**, **2C**). Together, these observations suggest that in the *rad-54.L* mutant, the inability to remove RAD-51 from DSBR sites prevents the stable association of subsequent factors such as MSH-5 to DSBR sites. However, in the *rad-54.B* mutant, the early steps of meiotic DSBR progress relatively normally.

**Fig 2.**
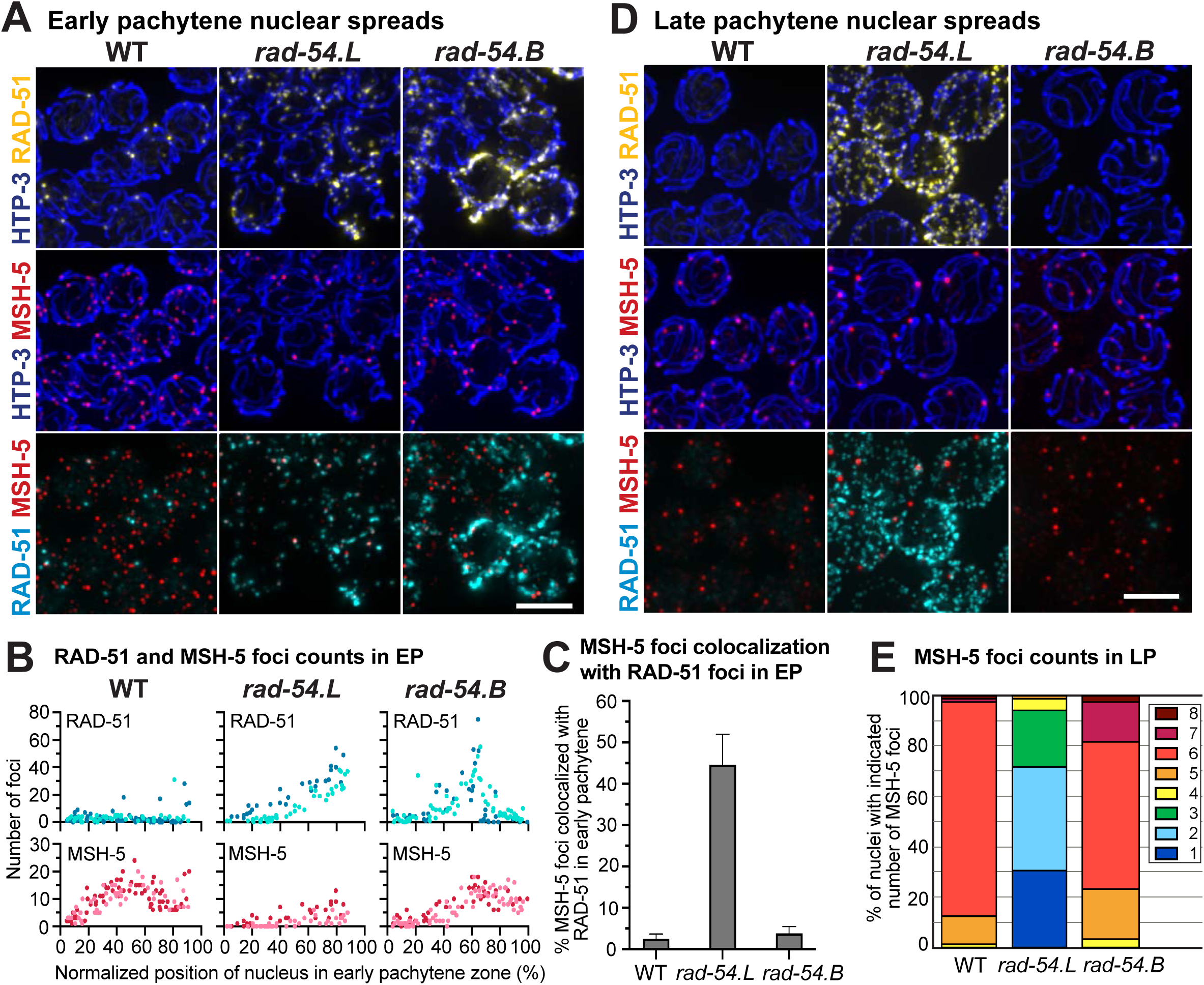
Differences in progression of meiotic recombination in *rad-54.L* and *rad-54.B* mutants.) Max-projected images of early pachytene nuclear spreads from WT, *rad-54.L(me98)*, and *rad-54.B(gk340656)* gonads, immunostained for RAD-51, meiotic recombination protein MSH-5 and chromosome axis protein HTP-3. Scale bar represents 5 μm. B) Quantification of RAD-51 and MSH-5 foci numbers in early pachytene nuclei from spread gonads. Each data point represents an individual nucleus, showing its relative position within the early pachytene zone and number of RAD-51 or MSH-5 foci. Light and dark colors in the same plot represent the two different gonads used in quantification. In the *rad-54.B* mutant, MSH-5 foci accumulate with some delay, but foci numbers plateau at WT-like levels (WT vs *rad-54.B*, first half of early pachytene zone, p<0.0001, second half of early pachytene zone, n.s). The *rad-54.L* mutant has reduced numbers of MSH-5 foci compared to WT and the *rad-54.B* mutant (WT vs *rad-54.L*, p<0.0001 for either half of the early pachytene zone, *rad-54.B* vs *rad-54.L*, p<0.0001 for either half of the early pachytene zone). Statistical significance was assessed using the Mann Whitney test. C) Percentage of MSH-5 foci colocalizing with RAD-51 foci in early pachytene spread nuclei (total numbers of MSH-5 foci analyzed: WT, n=1083; *rad-54.L*, n=177; *rad-54.B*, n=808). Error bars indicate 95% confidence interval. D) Max-projected images of RAD-51, MSH-5 and HTP-3 immunostaining in late pachytene spread nuclei, illustrating approximately normal numbers of MSH-5 foci in the *rad-54.B* mutant and persistence of high levels of RAD-51 foci and reduced numbers of MSH-5 foci in the *rad-54.L* mutant. Scale bar represents 5 μ m. E) Quantification of MSH-5 foci in late pachytene nuclear spreads.

Consistent with the successful progression of meiotic recombination and largely normal designation and formation of COs, MSH-5 foci counts in late pachytene spreads from the *rad-54.B* mutant were comparable to those observed in WT (**Fig 2D**, **2E**). This contrasts with the *rad-54.L* mutant, where late pachytene nuclei retained abundant hyperaccumulated RAD-51 foci and MSH-5 foci occurred in reduced numbers (**Fig 2D**, **2E**). These findings further support the conclusion that *rad-54.B* mutants are largely proficient for progression of meiotic DSBR, while *rad-54.L* mutants are severely compromised for progression of DSBR beyond the formation of early RAD-51 bound intermediates.

### RAD-51 hyperaccumulation in *rad-54.B* mutants is regulated by CHK-2 activity

As hyperaccumulation of RAD-51 foci in *rad-54.B* mutants is confined to a limited region of the germline similar to where RAD-51 foci are observed in WT germlines, we investigated the relationship of this hyperaccumulation phenotype to the activity of protein kinase CHK-2, a key regulator of multiple events in *C. elegans* meiosis.

CHK-2 activity is turned on at meiotic prophase onset, and it is essential for nuclear reorganization at meiotic entry, homolog pairing, normal SC assembly, and the formation of programmed meiotic DSBs (Kim et al., 2015; MacQueen and Villeneuve, 2001; Oishi et al., 2001; Rosu et al., 2013; Stamper et al., 2013). Further, CHK-2 activity is turned off at the transition from early to late pachytene, when the requirements for the “CO assurance checkpoint”—that every chromosome has a DSBR intermediate that is eligible to be differentiated into a CO—are fulfilled, as evidenced by the observation of an extended “CHK-2 active zone” in mutants that fail to form CO intermediates (Kim et al., 2015; Rosu et al., 2013; Stamper et al., 2013; Woglar et al., 2013). At this transition from early to late pachytene, several key changes occur in the meiotic program, such as the shutting down of the DSB-induction machinery, changes in the molecular requirements for DSBR, and a switch in the preferred DSBR repair template from homologous chromosome to sister chromatid (Hayashi et al., 2007; Rosu et al., 2013, 2011; Stamper et al., 2013).

Visualization of SUN-1 pS24 as a marker of CHK-2 activity (Penkner et al., 2009; Woglar et al., 2013) allowed us to make several inferences regarding RAD-51 hyperaccumulation and meiotic prophase progression in the *rad-54.B* mutant. First, co-staining for SUN-1 pS24 and RAD-51 revealed a striking temporal/spatial correspondence between the abrupt decline of RAD-51 hyperaccumulation and the end of the CHK-2 active zone (**Fig 3A**). The rapid disappearance of RAD-51 immediately follows the shut-down of CHK-2 activity; this suggests that hyperaccumulation of RAD-51 in the *rad-54.B* mutant may depend on CHK-2 activity. Further, “outlier” nuclei with high RAD-51 signal in the late pachytene region of the gonad also consistently exhibited SUN-1 pS24 signal, strengthening the inference that CHK-2 activity state and RAD-51 hyperaccumulation are linked. Second, quantification of the length of the CHK-2 active zone did not reveal any difference between WT and the *rad-54.B* mutant (**Fig 3A**, **3B**), indicating apparently normal timing of progression from early pachytene to late pachytene in the *rad-54.B* mutant. This implies that the requirements of the CO assurance checkpoint are being satisfied in a timely manner, consistent with our observations regarding successful progression of meiotic DSBR.

**Fig 3.**
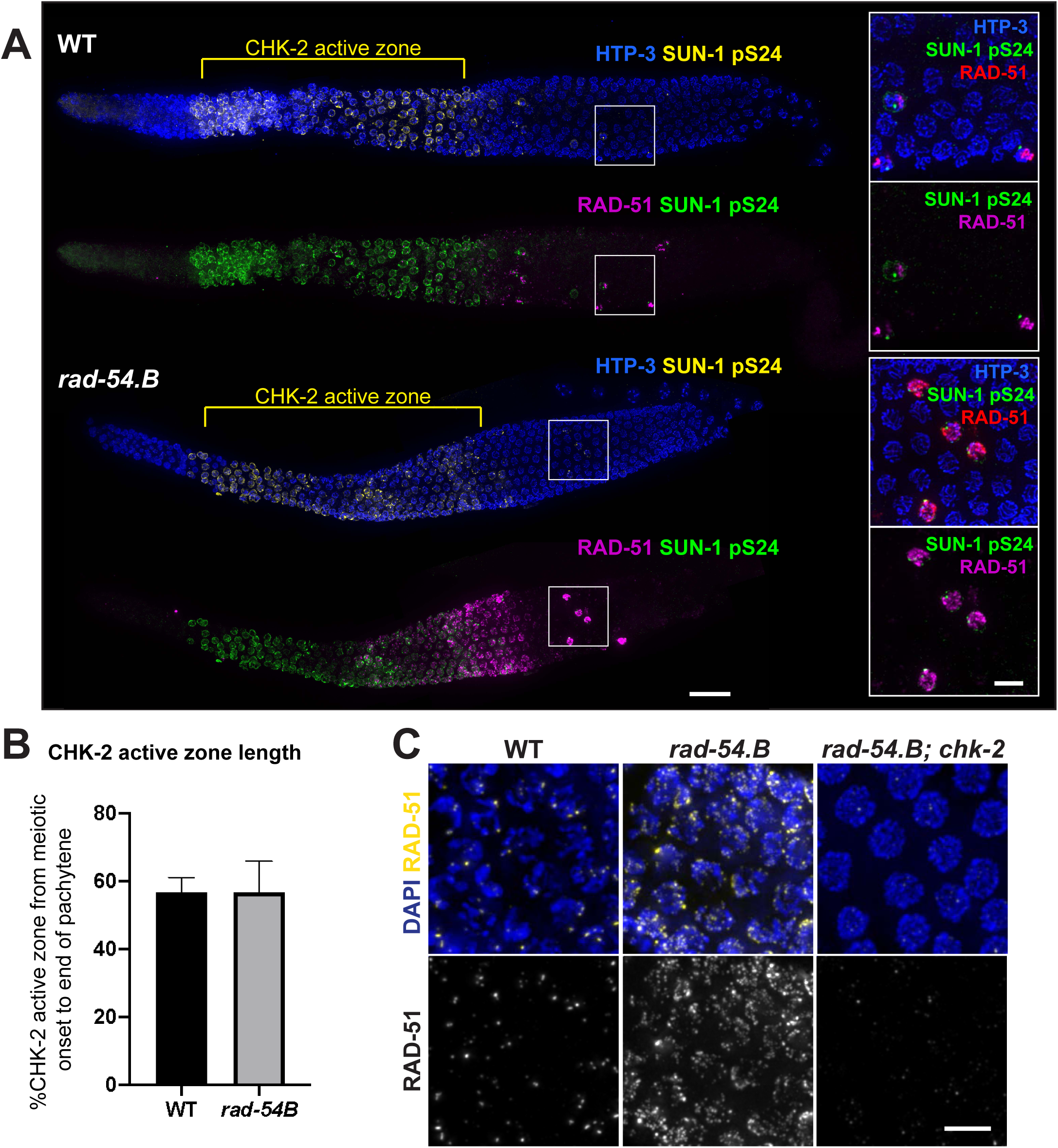
Relationship between CHK-2 activity and RAD-51 hyperaccumulation in the *rad-54.B* mutant. A) Max-projected images of whole-mount gonads stained for HTP-3, RAD-51, and SUN-1 pS24, an indicator of activity of protein kinase CHK-2, showing comparable lengths of the “CHK-2 active zones” in WT and *rad-54.B(gk340656)* gonads and illustrating that an abrupt drop in hyperaccumulated RAD-51 in the *rad-54.B* mutant coincides with the decline in CHK-2 activity that marks the transition from early to late pachytene. Scale bar represents 20 µm. Insets on the right show zoomed-in fields of nuclei from the late pachytene regions that include a few “outlier nuclei” with high levels of both RAD-51 and SUN-1 pS24, reflecting the normal operation of meiotic checkpoints in the *rad-54.B* mutant. Scale bar in insets represents 5 µm. B) Quantification of length of the CHK-2 active zone, defined as the contiguous region where the majority of nuclei in each cell row exhibited SUN-1 pS24 staining. Error bars represent standard deviation. Numbers of gonads analyzed: WT, n=18; *rad-54.B*, n=20. C) Max-projected images of RAD-51 staining in whole-mount gonads. Depicted nuclei are at the early pachytene stage for WT and *rad-54.B* (full genotype: *rad-54.B; egl-1 yIs34 oxTi633*) or from the equivalent position in the *rad-54.B; chk-2* gonad (full genotype: *rad-54.B; egl-1 chk-2*). Scale bar represents 5 µm.

Consistent with our hypothesis that CHK-2 activity is necessary for the hyperaccumulation of RAD-51 in the *rad-54.B* mutant, we found that RAD-51 hyperaccumulation was abrogated in a *rad-54.B; chk-2* double mutant (**Fig 3C, S3**). As *chk-2* mutants lack meiotic DSBs, however, this experiment did not distinguish whether RAD-51 hyperaccumulation is dependent on CHK-2 activity *per se* or on DSB formation; this issue will be addressed below.

### RAD-51 accumulates at DSBR sites in *rad-54.L* mutant germ cells but accumulates on unbroken DNA in the *rad-54.B* mutant

The observed differences in the dynamics of accumulation of RAD-51 foci and progression of DSBR in *rad-54.L* and *rad-54.B* germ lines raised the possibility that the way in which hyperaccumulated RAD-51 associates with chromosomes might be fundamentally different between these mutants. To address this possibility, we used structured illumination microscopy (SIM) to examine the morphology and spatial organization of RAD-51 foci in spread nuclei immunostained for RAD-51 (**Fig 4A**). SIM imaging of early pachytene nuclear spreads showed that while *rad-54.L* nuclei have an elevated number of RAD-51 foci, the RAD-51 foci have several similarities to those seen in WT. First, similarly to meiotic DSBR foci in WT meiosis (Woglar and Villeneuve, 2018), RAD-51 foci in the *rad-54.L* mutant are usually associated with the chromosome axes. Second, in WT nuclei, RAD-51 foci are often observed as elongated foci or doublets, which are interpreted to represent RAD-51 localizing on both resected ends of a DSB (Woglar and Villeneuve, 2018); RAD-51 foci in the *rad-54.L* mutant are similarly observed as extended singlets or doublets, with some foci exhibiting hyper-elongation. In *rad-54.B* nuclei, in contrast, RAD-51 foci are often not associated with the chromosome axis, and foci exhibit more variability in morphology. These data, together with the differences in DSBR progression in *rad-54.L* and *rad-54.B* mutants, suggested that the hyperaccumulated RAD-51 in these two *rad-54* family mutants might represent RAD-51 localizing at different underlying DNA structures.

**Fig 4.**
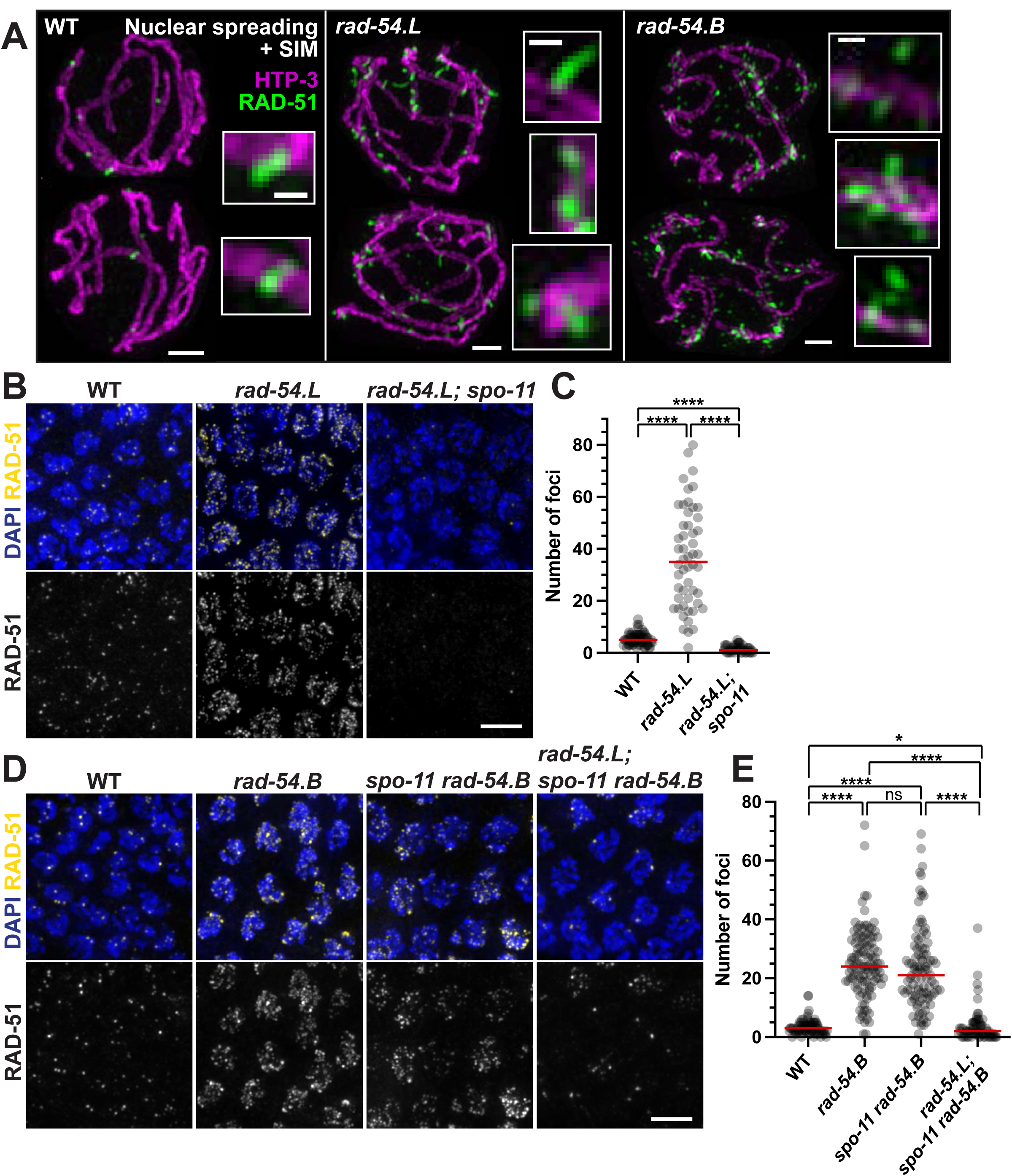
Evidence supporting RAD-51 hyperaccumulation at unbroken DNA in *rad-54.B* mutant. A) Max-projected images of RAD-51 and axis marker HTP-3 in spread early pachytene nuclei imaged by structured illumination microscopy (SIM), illustrating differences in the appearance of hyperaccumulated RAD-51 in *rad-54.L(me98)* and *rad-54.B(gk340656)* mutants (see main text). Scale bars for whole nuclei represent 1 µm; scale bars for zoomed-in insets represent 200 nm. B) Representative fields of nuclei from max-projected images of whole-mount gonads, taken from the zones of maximum accumulation of RAD-51 foci; scale bar represents 5 µm. C) Quantification of RAD-51 foci for the genotypes shown in panel B. RAD-51 foci counts were conducted in the zone of maximum accumulation of RAD-51 foci for each genotype. Each circle represents a nucleus; red lines indicate median values. Numbers of nuclei analyzed (n) and median numbers of foci (m) were as follows: WT, n=66, m=5; *rad-54.L*, n=52, m=35; *rad-54.L; spo-11*, n=49, m=1. Statistical significance was assessed with a Mann Whitney test; ****, p<0.0001. D) Representative fields of nuclei from max-projected images of whole-mount gonads, taken from the zones of maximum accumulation of RAD-51 foci. Scale bar represents 5 µm. E) Quantification of RAD-51 foci for the genotypes shown in (D). WT, n=81, m=3; *rad-54.B*, n=115, m=24; *spo-11 rad-54.B*, n=99, m=21; *rad-54.L; spo-11 rad-54.B*, n=71, m=2. Statistical significance was assessed with a Mann Whitney test; ns, p>0.05; *, p<0.05; ****, p<0.0001.

Consistent with this hypothesis, we found that the *rad-54.L* and *rad-54.B* mutants differ regarding whether hyperaccumulation of RAD-51 is dependent on the formation of meiotic DSBs. We confirmed previous work showing that hyperaccumulation of RAD-51 foci is abolished in a *rad-54.L; spo-11* double mutant, which lacks the enzyme responsible for generating meiotic DSBs (Dernburg et al., 1998; Mets and Meyer, 2009) (**Fig 4B**, **4C**, **S4A**). This indicates that RAD-51 hyperaccumulation in the absence of RAD-54.L is DSB-dependent and likely occurs at meiotic DSBR sites. In contrast, hyperaccumulation of RAD-51 foci in the absence of RAD-54.B occurs independently of meiotic DSBs, as hyperaccumulation persisted in the *spo-11 rad-54.B* double mutant (**Fig 4D**, **4E**, **S4B**). We infer that loss of *rad-54.B* function results in hyperaccumulation of RAD-51 on unbroken DNA, similarly to the previously-described DSB-independent accumulation of meiotic recombinase Dmc1 in *S. cerevisiae* mutants lacking the RAD-54.B ortholog Rdh54/Tid1 (Holzen et al., 2006).

The striking difference between the *rad-54.L* and *rad-54.B* mutants regarding DSB-dependence/independence of RAD-51 hyperaccumulation dovetails with our data showing that meiotic DSBR is stalled at an early intermediate in the *rad-54.L* mutant, but progresses successfully in the *rad-54.B* mutant. Together these findings indicate a major role for RAD-54.L in promoting progression of meiotic DSBR through a mechanism involving the removal of RAD-51 from early recombination intermediates, and a distinct role for RAD-54.B in preventing the accumulation of RAD-51 on unbroken DNA. Further, the demonstration of DSB-independent RAD-51 hyperaccumulation in the *spo-11 rad-54.B* double mutant indicates that loss of hyperaccumulation in the *rad-54.B; chk-*2 double mutant is not due to the lack of programmed DSBs, but instead reflects a role for CHK-2 in enabling association of RAD-51 with unbroken DNA when RAD-54.B is absent (**S3**).

Surprisingly, examination of RAD-51 localization in a *rad-54.L; spo-11 rad-54.B* triple mutant revealed that RAD-51 hyperaccumulation was greatly attenuated compared with the *spo-11 rad-54.B* double mutant (**Fig 4D**, **4E**, **S4B**). Specifically, while *rad-54.L; spo-11 rad-54.B* triple mutant germ lines had a few “outlier” nuclei with high levels of RAD-51 foci, the majority of nuclei had zero or only one or two RAD-51 foci with peak intensities above a baseline threshold for confident focus calling. Moreover, the 1-2 bright foci detected in most meiotic prophase nuclei likely represent the persistence of RAD-51 accumulated at sites of DNA damage incurred during DNA replication in the absence of both RAD-54.L and RAD-54.B, as such foci are also present in the mitotic proliferation zone at the distal end of the germ line in both *rad-54.L; rad-54.B* and *rad-54.L; spo-11 rad-54.B* mutant gonads (**Fig S4**). The unexpected requirement for RAD-54.L to achieve DSB-independent RAD-51 hyperaccumulation suggests that in the absence of RAD-54.B, RAD-54.L promotes promiscuous binding of RAD-51 to unbroken dsDNA.

To further investigate the contribution of RAD-54.L to promoting DSB-<u>independent</u> RAD-51 hyperaccumulation, we created a *rad-54.L* missense allele (*rad-54.L(me177)*, referred to as *rad-54.L(K238R)*) encoding a predicted ATPase-dead version of the RAD-54.L protein (Clever et al., 1999; Petukhova et al., 1999). *rad-54.L(K238R)* single mutant gonads exhibit persistent hyperaccumulation of RAD-51 comparable to that observed in the *rad-54.L(me98)* null mutant (hereafter referred to as *rad-54.L(null)*) (**S5**), consistent with the ATP-dependent motor activity of RAD-54.L being required to promote removal of RAD-51 at meiotic DSBR sites. Immunostaining of *rad-54.L(K238R); spo-11 rad-54.B* germ lines (**S6, S7**) further revealed a substantial attenuation of SPO-11-independent RAD-51 hyperaccumulation relative to *spo-11 rad-54.B*. However, the residual RAD-51 immunostaining observed in *rad-54.L(K238R); spo-11 rad-54.B* germ lines was also distinguishable from that observed in *rad-54.L(null); spo-11 rad-54.B* germ lines; whereas 1-2 bright RAD-51 foci were detected in most nuclei for both genotypes (presumably reflecting persistence of replication-associated DNA damage as discussed above), additional residual foci were more abundant in *rad-54.L(K238R); spo-11 rad-54.B* germ lines. This intermediate abundance of RAD-51 foci in the ATPase-dead mutant background suggests that RAD-54.L may promote the association of RAD-51 with unbroken DNA through both ATPase-dependent and ATPase-independent mechanisms.

### Extensive colocalization of RAD-54.L with both normal and hyperaccumulated RAD-51 foci

To better understand the roles of RAD-54.L and RAD-54.B, we investigated their localization in meiotic prophase germ cells. We assayed RAD-54.L localization by immunostaining for RAD-54.L::YFP in the germ lines of worms expressing a functional *rad-54.L::YFP* transgene in a *rad-54.L* null background (Stergiou et al., 2011). In whole-mount gonad preparations, RAD-54.L::YFP immunostaining was detected predominantly in the nucleoplasm (and variably in the nucleolus) (**Fig 5A-B**, **S8**). Thus, to visualize potential chromosome-associated RAD-54.L::YFP signals, we used a nuclear spread preparation to release nucleoplasmic protein pools. Nuclear spreading revealed RAD-54.L::YFP foci in early pachytene nuclei that exhibited essentially complete colocalization with RAD-51 foci, in both WT and *rad-54.B* mutant backgrounds (**Fig 5C**, **5D**). This suggests that RAD-54.L colocalizes with chromosome-associated RAD-51 regardless of whether RAD-51 is bound at DSBR sites or at unbroken DNA. Further, SIM imaging of these RAD-54.L::YFP foci showed that RAD-54.L::YFP foci are associated with chromosome axes and often occur as doublets or elongated singlets, similar to and colocalizing with RAD-51 (**Fig 5E**). In highly spread nuclei, partial spatial separation of RAD-51 and RAD-54.L::YFP signals became evident (**Fig 5E, bottom**), suggesting that RAD-51 and RAD-54.L associate with adjacent but non-identical portions of the underlying DNA molecules. However, we could not discern a stereotyped configuration of RAD-54.L::YFP relative to RAD-51 at these sites, as a variety of colocalization patterns were observed.

**Fig 5.**
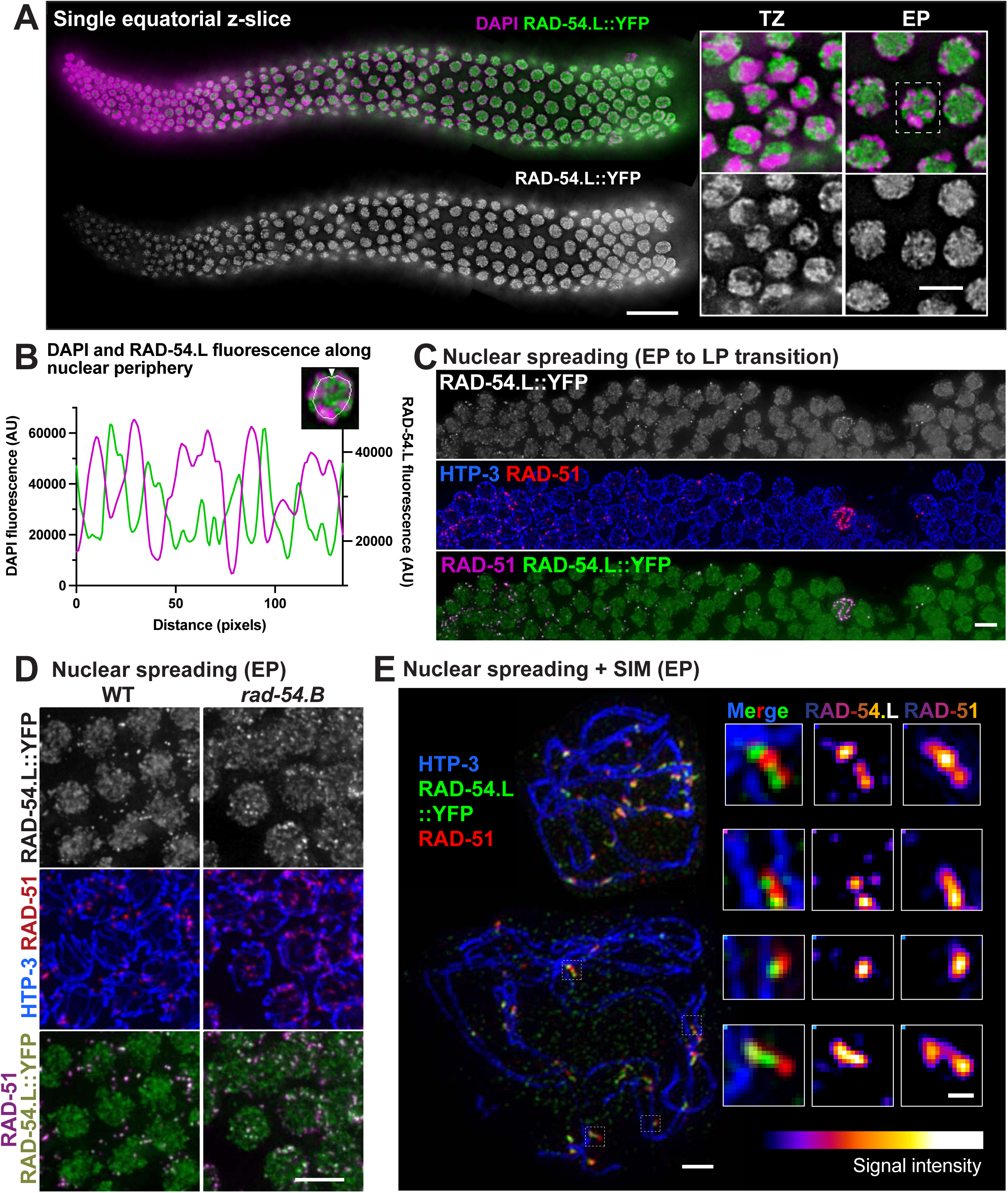
Localization of RAD-54.L. A) Image of whole-mount gonad from a *rad-54.L(null); opIs257[rad-54.Lp::rad-54.L::YFP::rad-54.L 3’UTR + unc-119(+)]* worm immunostained for RAD-54.L::YFP. The image represents a single z-slice showing an equatorial view of nuclei (instead of a max-projection). Scale bar represents 20 µm. Insets at right depict zoomed-in fields of nuclei from the transition zone (TZ) and early pachytene zone (EP), illustrating that the preponderance of RAD-54.L::YFP detected in whole-mount nuclei is not associated with the DAPI-stained chromatin. The scale bar for insets represents 5 µm. B) Quantification of fluorescence levels for DAPI (magenta) and RAD-54.L::YFP (green) along the periphery of the depicted nucleus; image represents a single equatorial Z-slice. Fluorescence was measured along the line indicated, starting at the arrowhead on the top and going in a counter-clockwise direction. This example nucleus is also indicated by a dotted box in (A). C) Max-projected representative field of view depicting the early pachytene (EP) to late pachytene (LP) transition in a spread gonad; scale bar indicates 5 µm. Nuclear spreading reveals a subset of RAD-54.L::YFP localizing in foci that colocalize with RAD-51. D) Max-projected images of immunostained spread early pachytene nuclei, illustrating extensive colocalization of RAD-51 foci with RAD-54.L::YFP foci in both WT (*rad-54.L; opIs257*) and *rad-54.B* (*rad-54.L; rad-54.B(gk340656); opIs257*) backgrounds. Scale bar indicates 5 µm. E) Max-projected early pachytene chromosome spreads imaged with SIM. Shown on the left side are two nuclei with different degrees of spreading; scale bar indicates 1 µm. Insets on the right are zoomed-in images of instances of RAD-51 and RAD-54.L::YFP colocalization from the widely-spread bottom nucleus. Fluorescence intensity of RAD-54.L and RAD-51 are depicted with an color look-up table (“LUT Fire” in ImageJ). Scale bar represents 200 nm.

### Localization of RAD-54.B at a subset of meiotic DSBR sites

To visualize RAD-54.B, we used CRISPR-Cas9 genome editing to generate a strain expressing RAD-54.B::GFP from the endogenous *rad-54.B* locus. In whole-mount gonads, RAD-54.B::GFP was detected predominantly in the nucleoplasm and the nucleolus, as observed for RAD-54.L::YFP (**Fig 6A**, **6B**). Further, nuclear spreading likewise revealed a subpopulation of RAD-54.B::GFP in pachytene nuclei localizing to foci associated with the chromosome axis (**Fig 6C-D**). While a subset of RAD-54.B::GFP foci colocalized with RAD-51 foci, however, the majority did not; conversely, the majority of RAD-51 foci did not colocalize with RAD-54.B::GFP foci (**Fig 6C**, **6E**). This contrasts sharply with the essentially complete coincidence observed for RAD-51 and RAD-54.L::YFP foci (above), presumably reflecting distinct roles and contributions of RAD-54.L and RAD-54.B.

**Fig 6.**
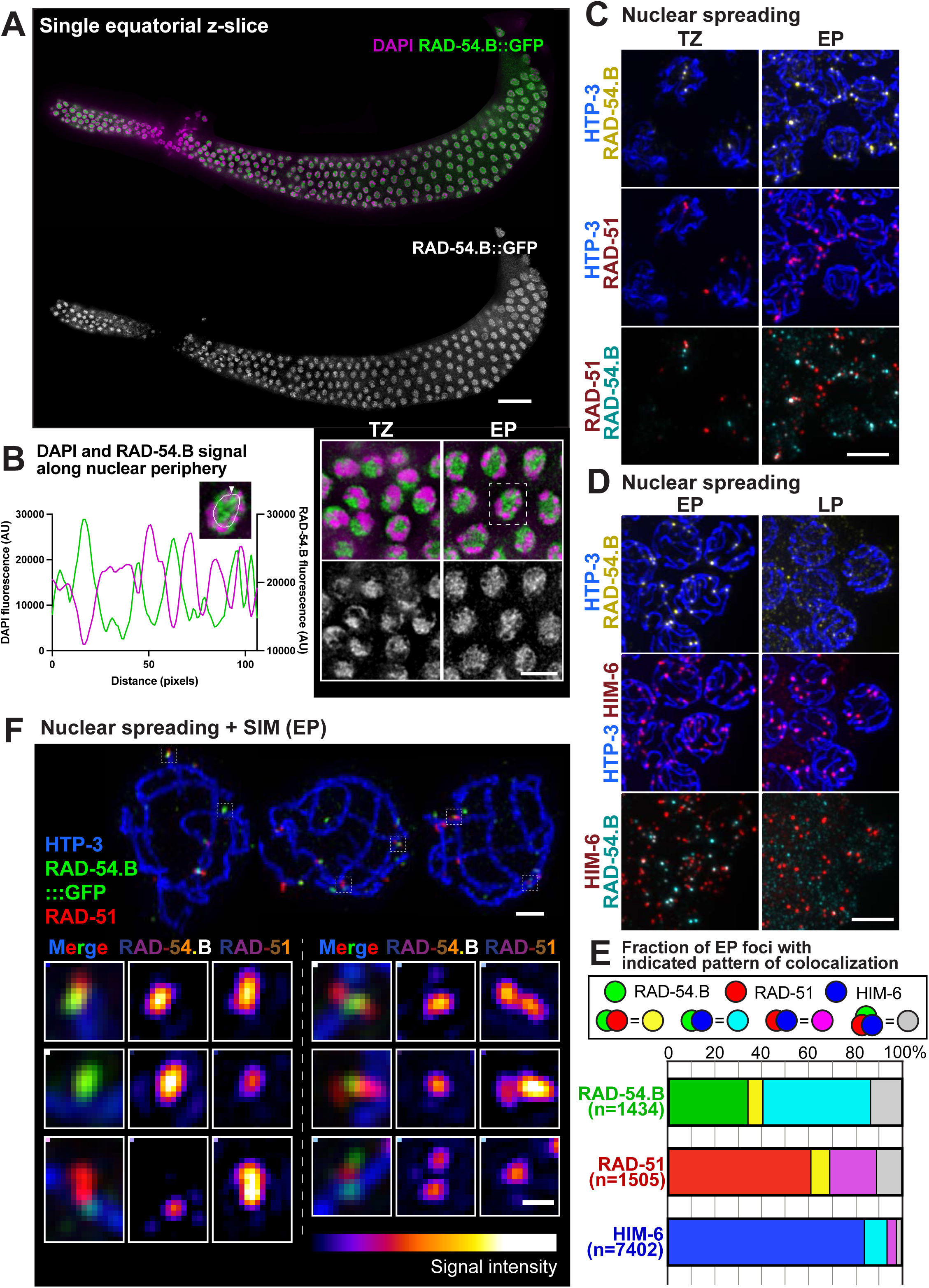
Localization of RAD-54.B. A) Image of whole-mount *rad-54.B(gt3402[rad-54.B::GFP])* gonad immunostained for RAD-54.B::GFP and DAPI. The image represents a single z-slice showing an equatorial view of nuclei (instead of a max-projection). Scale bar represents 20 µm. Insets at right-bottom depict a zoomed-in field of nuclei from the transition zone (TZ) and early pachytene zone (EP), showing that similarly to RAD-54.L, the preponderance of RAD-54.B is not associated with DAPI-stained chromatin. The scale bar for insets represents 5 µm. B) Quantification of fluorescence levels of DAPI (magenta) and RAD-54.B::GFP (green) along the periphery of the depicted nucleus, as in Fig 5. This example nucleus is also indicated by a dotted box in (A). C) Max-projected images of RAD-54.B::GFP and RAD-51 immunostaining in spread nuclei from the transition zone (TZ) and the early pachytene (EP) zone. Release of soluble RAD-54.B by nuclear spreading reveals distinct axis-associated RAD-54.B foci, a fraction of which colocalize with RAD-51 foci. Scale bar represents 5 µm. D) Max-projected images of RAD-54.B::GFP and HIM-6 immunostaining in spread nuclei in early pachytene (EP) and late pachytene (LP). Distinct axis-associated RAD-54.B::GFP foci detected early pachytene frequently colocalize with HIM-6 foci; axis-associated RAD-54.B::GFP signals diminish in late pachytene. Scale bar represents 5 µm. E) Quantification of colocalization between RAD-54.B::GFP, RAD-51, and HIM-6 foci in early pachytene chromosome spreads. For each species of focus, the fractions of solo foci and foci colocalizing with one or both of the other proteins are indicated. F) Max-projected early pachytene chromosome spreads imaged with SIM; scale bar represents 1 µm. Insets on bottom are zoomed-in images of instances of colocalization of RAD-51 and RAD-54.B::GFP. Fluorescence intensity of RAD-54.B and RAD-51 are depicted with an color look-up table (“LUT Fire” in ImageJ). Scale bar in insets represents 200 nm.

Examination of sites of RAD-51 and RAD-54.B::GFP colocalization by SIM imaging (**Fig 6F**) revealed overlapping or adjacent but non-identical localization of the two proteins at these sites, but no discernable stereotypical spatial relationship was evident. We observed cases of a RAD-51 singlet adjacent to a RAD-54.B::GFP singlet, as well as cases in which a RAD-51 doublet flanks RAD-54.B::GFP, or vice versa.

We further assessed the relationship of RAD-54.B::GFP foci to meiotic DSBR sites by costaining for HIM-6::HA (BLM helicase). This analysis revealed: a) partial colocalization between RAD-54.B::GFP foci and HIM-6::HA foci in early pachytene nuclei, and b) lost or diminished axis-associated RAD-54.B::GFP signal in late pachytene nuclei, in which HIM-6::HA localization is restricted to a single CO-designated site per chromosome pair (**Fig 6D**, **6E**).

We quantified colocalization between RAD-54.B::GFP, RAD-51, and HIM-6::HA in early pachytene spread nuclei co-stained for all 3 proteins (**Fig 6E**). For each protein, we quantified the fraction of foci that were observed alone or colocalizing with one or both other proteins. This analysis indicated that: a) the majority of RAD-54.B::GFP foci colocalize with RAD-51 (92/1434, 6.4%), HIM-6::HA (660/1434, 46.0%), or both (195/1434, 13.6%), but b) only a minority of RAD-51 foci (287/1505, 19.1%) or HIM-6 foci (889/7402, 12.0%) colocalize with RAD-54.B::GFP (**Fig 6E**). Colocalization of RAD-54.B foci with foci marking meiotic DSBR sites suggests that in addition to its role in antagonizing accumulation of RAD-51 on unbroken DNA, RAD-54.B may also play a role (albeit non-essential) in meiotic DSB repair.

In addition to the colocalization analysis, the preservation of temporal/spatial organization of nuclei in the germ line in the imaged samples allowed us to plot the positions of the different classes of foci (categorized by colocalization pattern) along an axis corresponding to temporal progression through the early pachytene stage (**S9A**). While all types of foci were present throughout most of this spatial “time course”, analysis of relative distributions and median positions showed that RAD-51 foci tend to appear earlier than RAD-54.B::GFP foci, which in turn tend to precede HIM-6::HA foci.

Further, HIM-6::HA foci that colocalize with RAD-51 (with or without RAD-54.B::GFP) tend to precede HIM-6::HA foci that colocalize only with RAD-54.B::GFP, and these further tend to precede HIM-6::HA foci that are observed on their own. Our combined colocalization and temporal analyses suggest that RAD-54.B may associate transiently with DSBR intermediates during their transition from early intermediates marked by RAD-51 alone into later post-strand-exchange intermediates marked by HIM-6 alone.

Although the majority of RAD-54.B::GFP foci colocalized with RAD-51 and/or HIM-6::HA foci, suggesting participation in DSBR, 487/1434 (34%) of RAD-54.B::GFP foci did not colocalize with either. Interestingly, these solo RAD-54.B::GFP foci tended to be dimmer and appear later than RAD-54.B::GFP foci colocalizing with RAD-51 and/or HIM-6::HA (**S9A, S9B**). We speculate that these later, dimmer solo RAD-54.B::GFP may represent RAD-54.B associated with unbroken DNA.

### RAD-54.B contributes to successful completion of meiosis in a *rad-54.L* partial loss-of-function background

Although our initial analyses had demonstrated that RAD-54.B is largely dispensable for successful meiosis, localization of RAD-54.B in foci at DSBR sites during meiotic prophase raised the possibility that RAD-54.B might nevertheless contribute in a non-essential way to meiotic DSB repair. We sought evidence for participation of RAD-54.B in meiotic DSBR in two ways.

We first used a genetic assay to assess the distribution of COs relative to a set of Chromosome V SNP markers in control and *rad-54.B* mutant backgrounds (**S10**). Consistent with our cytological observation of 6 COSA-1 foci in late pachytene nuclei in *rad-54.B* mutants, we did not detect a reproducible difference between control and the *rad-54.*B mutant in the total frequency of Chromosome V COs. However, data from two independent experiments did suggest that the distribution of COs among intervals is modestly altered in a *rad-54.B* mutant background (Expt. 1, p=0.03; Expt. 2, p=0.003).

More definitive evidence for a contribution of RAD-54.B to meiotic DSBR was provided by experiments using *rad-54.L(me139),* a partial loss-of-function mutant (Akerib et al., 2022), as a sensitized genetic background. Assessment of the viability of embryos produced by WT, *rad-54.B* and *rad-54.L(me139)* single mutants, and the *rad-54.L(me139); rad-54.B* double mutant revealed that loss of RAD-54.B in the *rad-54.L(me139)* mutant background resulted in a substantial decrease in embryonic viability (**Table 1**). In contrast to the 98% and 26% egg hatching rates observed for the *rad-54.B* and *rad-54.L(me139)* single mutants, respectively, the *rad-54.L(me139); rad-54.B* double mutant exhibited 100% embryonic lethality (0% egg hatching), similarly to a *rad-54.L* null mutant.

**Table 1:** RAD-54.B can contribute to successful reproduction and development in a *rad-54.L* partial loss-of-function background.

| Genotype | Total number of eggs | % Eggs hatching | % Eggs reaching adulthood | % Males (/adults) |
| --- | --- | --- | --- | --- |
| WT | 869 | 99.9 | 99.9 | 0 |
| *rad-54.B(gk340656)* | 996 | 98.2 | 97.9 | 0.2 |
| *rad-54.L(me139)*<sup>a</sup> | 961 | 26.3 | 15.2 | 15.8 |
| *rad-54.L(me139); rad-54.B(gk340656)* | 777 | 0 | 0 | NA |

For progeny viability counts, single L4 hermaphrodites (4 for each genotype) were placed on individual plates, then each worm was transferred to a new plate at 24 h intervals until 72 h post L4. Numbers of eggs and larvae were counted 24 h (time point 1) and 48 h (time point 2) after plating of the parent worm; numbers of adult hermaphrodites and males were counted 72-96 hours after plating (time point 3). “Total number of eggs” laid was taken to be the sum of eggs and larvae at either time point 1 or 2, or the total number of adult worms at time point 3, whichever was larger, to account for both undercounting of young larvae and larval lethality. % Eggs hatching = $100 \times (1 - \text{unhatched eggs at time point 2} / \text{Total eggs laid})$ .
<sup>a</sup> For *rad-54.L(me139)*, data are from (Akerib et al., 2022). The difference between “% Eggs hatching” and “% Eggs reaching adulthood” reflects larval arrest/lethality.

Further, assaying chromosome morphology in diakinesis oocytes with DAPI staining revealed that loss of *rad-54.B* function aggravates the diakinesis defects observed in the *rad-54.L(me139)* mutant background (**Fig. 7**). Six bivalents are usually observed in WT and *rad-54.B* diakinesis oocytes, reflecting successful formation of CO-based attachments between homologs and completion of DNA repair during earlier stages of prophase. In contrast, DAPI-stained bodies in diakinesis oocytes in the *rad-54.L(me139)* mutant exhibit more variability in size and shape, a phenotype indicative of a defect in DSBR, which may cause chromosome fragmentation and/or inappropriate associations (Alpi et al., 2003; Colaiácovo et al., 2003). This abnormal diakinesis phenotype was aggravated in the *rad-54.L(me139); rad-54.B* double mutant, where DAPI-stained bodies in diakinesis oocytes were also highly variable in size and shape and deviated even further from normal-appearing bivalents, frequently exhibiting a “stringy” appearance in which many DAPI bodies appeared to be connected by thin DAPI-stained threads or bridges. Further, the fraction of oocyte nuclei exhibiting abnormal phenotypes (*i.e.*, categorized as having fewer than 5 or greater than 6 DAPI bodies) was significantly higher in the *rad-54.L(me139); rad-54.B* double mutant than in the *rad-54.L(me139)* single mutant. Taken together, these data reveal a hidden capacity of RAD-54.B to contribute to the successful completion of meiotic DSBR.

**Fig 7.**
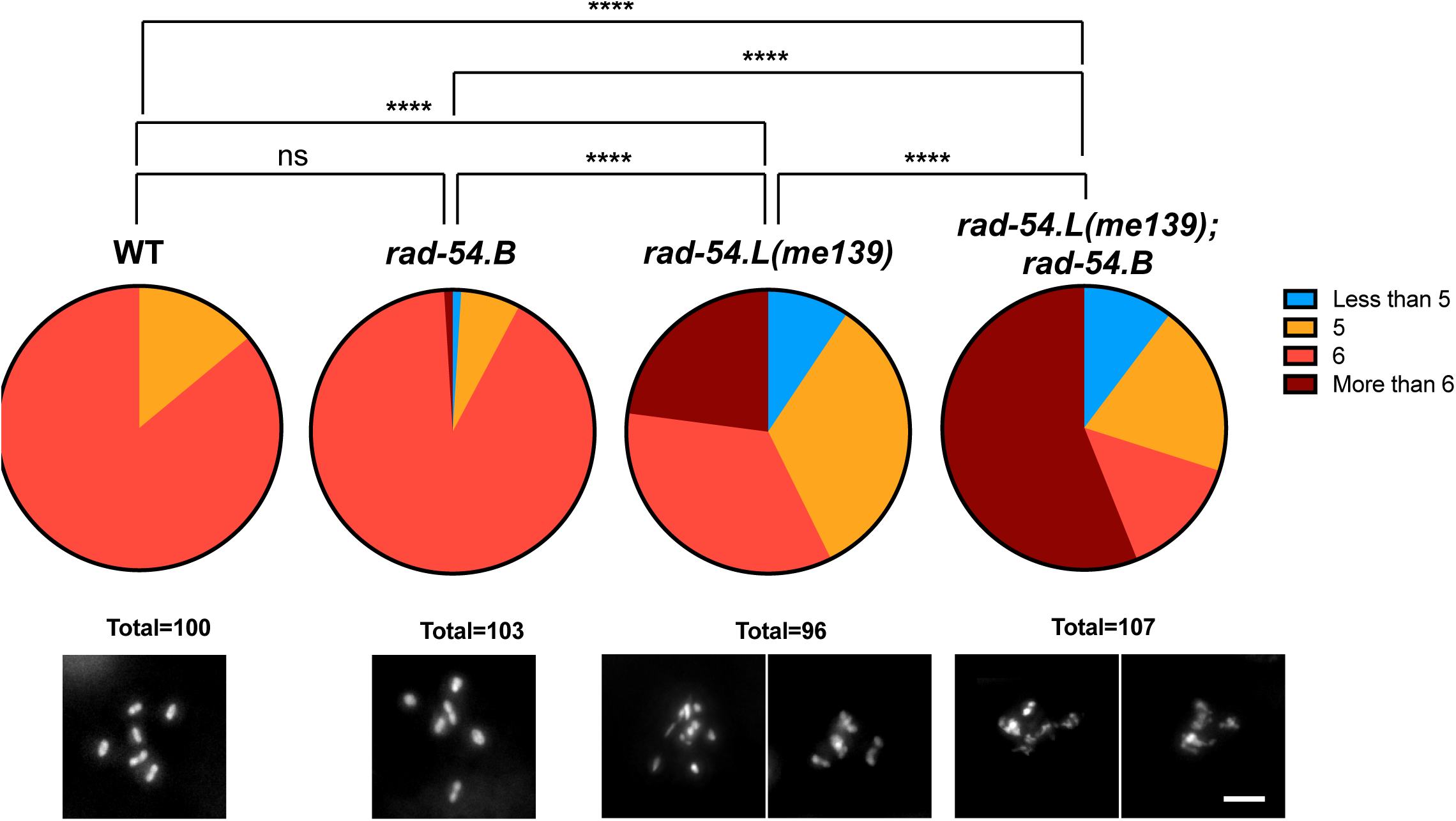
Evidence that RAD-54.B contributes to meiotic DSBR. A) Top, pie charts quantifying the fraction of oocytes displaying the indicated numbers of DAPI-stained bodies. Bottom, examples of DAPI-stained diakinesis oocytes for each genotype. *rad-54.L(me139); rad-54.B(gk340656)* oocytes had more irregularities in DAPI body morphology than *rad-54.L(me139)* oocytes, often appearing “stringy” and exhibiting more variability in size. While some *rad-54.L(me139)* and *rad-54.L(me139); rad-54.B* oocytes were scored as having 5-6 DAPI bodies, in many cases these did not represent WT-like bivalents, but instead may represent products of failed DSB repair such as chromosome fragments or fused chromosomes. Although DAPI-body number is an imperfect representation of such abnormal diakinesis figures, this quantification does capture the fact that the defective diakinesis phenotype caused by *rad-54.L(me139)* is aggravated by simultaneous loss of *rad-54.B*. Scale bar represents 5 μm. Statistical significance was assessed with a Mann Whitney test; ns, p>0.05; ****, p<0.0001.

## Discussion

Our current work clearly shows that RAD-54.L and RAD-54.B make very different contributions to the success of homologous recombination in the *C. elegans* germ line. We demonstrate a division of labor between and counterbalancing effects of these RAD-54 paralogs during *C. elegans* meiosis, with RAD-54.L in the crucial leading role and RAD-54.B as a supporting player.

We confirm prior work indicating an essential role for RAD-54.L in promoting the removal of RAD-51 from meiotic DSBR sites, and we extend this finding to demonstrate a requirement for a functional ATPase domain, implying that the translocase activity of RAD-54.L is important for its role in displacement of RAD-51. We further show that RAD-54.L is required for normal progression of subsequent steps in meiotic DSBR, including recruitment of the MutSg complex, consistent with RAD-51 removal and MutSg recruitment being mechanistically coupled events. Moreover, in the *rad-54.L* mutant background, RAD-51 colocalizes with MutSg at the small subset of DSBR sites where MutSg is recruited, reflecting impairment in the transition from recombinase-bound intermediates to post strand-exchange intermediates lacking RAD-51.

The strong mechanistic connections between RAD-51 and RAD-54.L are further reflected in their extensive colocalization, which suggests that RAD-51 filaments rarely occur without RAD-54.L being in close proximity, and conversely, that RAD-54.L rarely occurs in foci without RAD-51. RAD-54.L foci colocalized with RAD-51 foci may represent RAD-54.L translocases actively promoting homology search by mediating motor-driven one-dimensional movement of RAD-51-ssDNA filaments (Crickard et al., 2020b) and/or they may represent RAD-54.L translocases engaged in the process of RAD-51 removal, thereby driving strand exchange and D-loop formation (Crickard, 2021; Wright and Heyer, 2014). Alternatively, or in addition, colocalized foci may reflect a RAD-51-filament-stabilizing activity of RAD-54.L (see below).

In striking contrast to the central and essential role for RAD-54.L in meiotic DSBR, we found that RAD-54.B is largely dispensable for successful meiotic recombination. This indicates that despite *C. elegans* having only a single RAD-51 recombinase, the two RAD-54 paralogs must partner with RAD-51 in very different ways; further, RAD-54.B cannot substitute for RAD-54.L to complete the essential tasks of removing RAD-51 and enabling DSBR progression. However, localization of RAD-54.B at a minority of meiotic DSBR sites during wild-type meiosis and a modest delay in the appearance of MSH-5 foci in the *rad-54.B* mutant background together suggest that RAD-54.B likely does play a transient, albeit largely non-essential, role during meiotic DSBR. Based on prior studies identifying several disparate biochemical activities for RAD54 family translocases, we speculate that RAD-54.B might potentially function in multiple capacities at DSBR sites. For example, it might affect the structure of DSBR intermediates by modulating D-loop size or D-loop maturation, as proposed for yeast Rdh54/Tid1 (Keymakh et al., 2022; Shah et al., 2020), and/or it might modulate the activity of RAD-54.L in promoting RAD-51 removal. RAD-54.B functioning in such capacities at DSBR sites may underlie the modest difference in CO distribution observed in the *rad-54.B* mutant and the contribution of RAD-54.B to meiotic success in a *rad-54.L* partial loss-of-function background.

The most conspicuous function uncovered for RAD-54.B in this work is its role in preventing hyperaccumulation of RAD-51 on unbroken DNA. This finding parallels previous observations that the RAD-54.B ortholog Rdh54/Tid1 is required to inhibit the association of meiotic recombinase Dmc1 with unbroken DNA during *S. cerevisiae* meiosis (Holzen et al., 2006) and that both RAD-54.B and RAD-54.L orthologs contribute to antagonizing unproductive RAD51 accumulation and associated toxicity when RAD51 is overexpressed in vegetative yeast cells or cancer cells (Mason et al., 2015; Shah et al., 2010). The most straightforward interpretation is that the translocase activity of RAD54 proteins is responsible for removing recombinases from dsDNA. Such unproductive association of RAD51 or DMC1 with unbroken DNA has been proposed to reflect a relatively weak preference for ssDNA over dsDNA compared to bacterial RecA recombinases (Reitz et al., 2021). Thus, our observation that *C. elegans* RAD-51 exhibits a similar propensity for promiscuous association with unbroken DNA that is antagonized by RAD-54.B implies that this property is a conserved aspect of eukaryotic recombinases. However, our study also yielded a new and unexpected plot twist, namely that RAD-54.L itself is largely responsible for *<u>promoting</u>* the ectopic hyperaccumulation of RAD-51 on unbroken DNA when RAD-54.B is absent during *C. elegans* meiosis.

This unexpected finding that RAD-54.L promotes unproductive association of RAD-51 with unbroken DNA (in the *rad-54.B* mutant background) represents an important *in vivo* demonstration that a RAD54.L ortholog can act on RAD51 in opposing ways, *i.e.*, both to promote recombinase removal and to promote or stabilize recombinase association with DNA. While several studies have established that yeast Rad54 (RAD54.L ortholog) can stabilize Rad51 filaments *in vitro* (Mazin et al., 2003; Meir et al., 2022), there has been less evidence regarding the *in vivo* occurrence and relevance of filament-stabilizing activity. Thus, our observation that RAD-54.L is required for RAD-51 hyperaccumulation on unbroken DNA in the *rad-54.B* mutant helps to address this gap, providing evidence for its capacity to stabilize RAD-51 filaments *in vivo*. Interestingly, our analysis of the ATPase-dead allele, *rad-54.L(K238R)*, indicated that such RAD-51 promoting/stabilizing activity of RAD-54.L is partially dependent on RAD-54.L ATPase activity. We speculate that this reflects both ATPase-independent RAD-51-stabilizing activity, as well as ATPase-dependent RAD-51-promoting activity, perhaps involving translocation along and opening of dsDNA (Crickard et al., 2020b). Further, our data imply that opposing RAD-54.L activities can operate even within the same nucleus, with RAD-54.L promoting RAD-51 removal and progression of recombination at *bona fide* DSBR sites while simultaneously facilitating association of RAD-51 with unbroken DNA elsewhere in the nucleus. Importantly, however, our data clearly indicate that RAD-54.L is *<u>not</u>* required to promote/stabilize RAD-51 association with DNA at meiotic DSBR sites, as evidenced by the abundant DSB-dependent accumulation and persistence of RAD-51 in *rad-54.L* mutants.

How might progression through the meiotic program influence or be served by the observed disparate activities of RAD-54.L and division of labor between RAD-54.L and RAD-54.B? Our thinking in this regard is informed by our findings that: 1) the unproductive hyperaccumulation of RAD-51 at unbroken DNA in the *rad-54.B* mutant occurs in parallel with productive association of RAD-51 at meiotic recombination intermediates, and 2) RAD-51 hyperaccumulation occurs specifically in the “CHK-2 active zone” of the germ line and is dependent on CHK-2 activity. Several prior studies have shown that the window of activation of *C. elegans* CHK-2 from meiotic prophase onset through mid-pachytene corresponds to both the period of active DSB formation (Hinman et al., 2021; Rosu et al., 2013; Stamper et al., 2013) as well as the timing of engagement of a specialized “meiotic mode” of DSB repair involving modifications to the molecular requirements for processing and HR-mediated repair of DSBs (Hayashi et al., 2007; Rosu et al., 2011). We hypothesize that RAD-54.L (and/or RAD-51) may become hyperactivated within the CHK-2 active window as part of this meiotic mode of DSBR. We speculate that enhanced activity of such key recombination factors contemporaneously with DSB formation may be beneficial for promoting efficient homology search and strand exchange to enable both crossover formation and restoration of genome integrity. However, hyperactivation of RAD-54.L to augment DSBR may also increase its ability to stabilize or promote the formation of unproductive RAD-51 filaments on unbroken DNA, thereby necessitating the RAD-51-removal activity of RAD-54.B to antagonize this unproductive association.

This proposed hyperactivation of RAD-51 and/or RAD-54.L during *C. elegans* meiosis stands in striking contrast to the previously-described regulation of the orthologous proteins in *S. cerevisiae* meiotic cells, which actively down-regulate the strand-exchange activity of Rad51, in part through Rad54-phosphorylation-mediated inhibition of Rad54-Rad51 complex formation (Niu et al., 2009). However, these opposing modes of regulation of the RAD51 recombinase/RAD54L translocase partnership make sense in light of notable differences in the inventory of meiotic machinery components present in these two organisms. In *S. cerevisiae*, downregulation of Rad51 recombinase activity during meiotic prophase is important to enable interhomolog strand-exchange driven by the meiotic recombinase Dmc1 (in partnership with Rdh54). In contrast, *C. elegans* lacks DMC1, so RAD-51 is the sole recombinase available to create the crossovers needed to segregate homologous chromosomes and to promote HR-mediated repair of meiotic DSBs.

Together, the findings presented here contribute to a growing appreciation of RAD54.L and RAD54.B as highly versatile components in the recombination toolbox. Depending on the context in which they are deployed and how they are regulated, the diverse biochemical activities demonstrated for these RAD54 paralogs can serve to augment, modulate and/or counteract the activities of recombinases, thereby protecting genome integrity by ensuring repair outcomes that are achievable and appropriate for the situation at hand.

## Materials and Methods

### *C. elegans* Culturing, Genetics, and Gene Editing

Worms were grown using standard methods (Stiernagle, 2006) at 20°C, except for experiments depicted in **Fig 1D-E**, **Fig 2**, **Fig 3A-B**, **Fig 4A**, **Fig 6D, and S2**, where worms were grown at 22°C. *rad-54.B(gt3308)* was isolated in a screen for mutants with increased sensitivity to irradiation at the L1 larvae stage (González-Huici et al., 2017). *rad-54.B(gk340656)* was obtained from the Millions Mutations Project (Thompson et al., 2013). *rad-54.L(me177)*, *rad-54.B(gt3402[rad-54.B::GFP])*, and *rad-54.B(gt3328)* were generated using CRISPR-Cas9 gene editing using established methods (Dickinson et al., 2015, 2013; Dokshin et al., 2018; Paix et al., 2015; Wang et al., 2018; Ward, 2015); details are provided in Supplemental Information. A list of strains used in this study is provided in Supplemental Information.

### Immunofluorescence and Imaging

Immunofluorescence experiments using whole-mount gonads or spread nuclei were conducted as in (Martinez-Perez and Villeneuve, 2005; Pattabiraman et al., 2017; Woglar and Villeneuve, 2018); worms were dissected at 22-30 hours post L4. Details about antibodies used are available in Supplemental Information.

Image acquisition and processing were conducted as in (Roelens et al., 2019a; Woglar and Villeneuve, 2018). All images were acquired using a DeltaVision OMX Blaze microscope with a 100x NA 1.4 NA objective, with 200 nm spaced z-stacks for widefield images and 125 nm spaced z-stacks for SIM images. Images were processed using SoftWoRx and FIJI software (Preibisch et al., 2009); additional details are provided in Supplemental Information.

### Chromosome Tracing and Quantification of CO site Focus Distribution

Chromosome tracing and quantification of CO site distribution (**Fig 1D, S2**) were done manually using the SNT plugin on ImageJ (Arshadi et al., 2021). Gonad nuclear spreads prepared from AV695 and TG4252 worms were immunostained using antibodies for HTP-3, SYP-1, GFP, and MSH-5, and imaged using SIM. Nuclei in the late pachytene stage with 5 or 6 COSA-1::GFP and MSH-5 foci and with SYP-1 still localized along the entire lengths of the aligned homolog pairs were selected for quantification of CO site distribution. HTP-3 and SYP-1 were used as markers of the chromosome axis, while COSA-1::GFP and MSH-5 were used as markers for CO sites. Images were collapsed into single-channel 8-bit images and input into the ImageJ SNT plugin. For each selected nucleus, the lengths of all chromosomes were measured by manually tracing the chromosome using SNT, and the distance of each CO site to the nearest chromosome end was similarly quantified. *mnT12* fusion chromosomes and non-fusion chromosomes were distinguished after tracing by overall chromosome length, as *mnT12* chromosomes were longer than 15 µm, while all other chromosomes were shorter than 15 µm.

### Focus count quantification

Z-stack images of whole gonads were cropped to include only a single layer of germ cell nuclei. Nuclei that were well-separated from each other were manually segmented using HTP-3 signal in max-projections of these cropped images and polygon-shaped ROIs were saved in FIJI software. The FIJI 3D Maxima Finder plugin was used to identify foci within these ROIs and determine their peak brightness and the positions of their maxima (Ollion et al., 2013); details on parameters used are provided in Supplemental Information. From the 3D Maxima Finder output table of foci with their xyz peak positions and peak heights, foci were computationally assigned to ROIs corresponding to individual nuclei based on their positions. Nucleus-focus assignments were manually checked to remove overlap to generate the final focus counts. The xy position of each nucleus ROI was used to approximate its position within the region of quantification, *i.e.* the x-axis values in **Fig 1E** **and** **2B**.

For whole-gonad quantifications (**Fig 1E**), this analysis was performed across the entire gonad. For quantification in early pachytene nuclear spreads (**Fig 2B**, **2E**, **5E**, **S9**), the early pachytene zone was defined as starting at the cell row in which most nuclei have a RAD-51 focus and ending at the cell row in which most nuclei have 6 MSH-5 or HIM-6 foci. For quantification in late pachytene nuclear spreads (**Fig 2E**), the late pachytene zone was defined as starting at the cell row in which most nuclei have 6 MSH-5 foci and ending before the diplotene stage when HTP-3 axis staining loses its linear appearance. For both cases, the appropriate region was cropped and used for the quantification pipeline above. For quantification of RAD-51 foci in *rad-54.L* and *rad-54.B* mutants (**Fig 3B-E, S6**), a 10 cell-row wide zone corresponding to the region of peak accumulation of RAD-51 foci was manually identified and cropped before performing focus count quantification as described above.

### Colocalization Analysis

For colocalization analyses (**Fig 2C**, **6E**), the xyz positions and peak heights of foci were determined as outlined above. For each species of focus quantified (e.g. MSH-5 in **Fig 2C**), the distance from each focus to the nearest neighbor focus of the other species (e.g. RAD-51 in **Fig 2C**) was determined computationally. Foci were considered colocalized if the distance to the nearest neighbor was under the appropriate threshold, calculated as the smallest resolvable distance based on the numerical aperture of the objective (1.4) and wavelength (λ) used (0.61λ/1.4, *i.e.* 242 nm if λ=555 nm, 282 nm if λ=647 nm).

### DAPI Staining of Diakinesis Oocytes

Ethanol fixation and DAPI staining of diakinesis oocytes was conducted as in (Bessler et al., 2007).

### Meiotic crossover (CO) distribution assay using SNP mapping strategy

Meiotic CO distribution was assayed as described (Agostinho et al., 2013). Additional details are available in Supplemental Information.

## Acknowledgments

We are grateful to A. Dernburg and V. Jantsch for antibodies and the CGC, which is funded by NIH Office of Research Infrastructure Programs (P40 OD010440), for strains. We thank J. Mulholland, K. Lee, A. Kahn, C. Akerib, B. Roelens, C. Choi, and C. Uebel for technical assistance and discussions. This work was funded by an American Cancer Society Research Professor Award (RP-15-209-01-DDC) and NIH grant R35GM126964 to AMV, a Stanford Graduate Fellowship to KY, grant 1S10OD01227601 from the National Center for Research Resources (NCRR) to the Stanford Cell Sciences Imaging Facility, a Natural Science Foundation grant of Guangxi Zhuang Autonomous Region (2022GXNSFAA035435) to BW. The AG lab is supported by Korean taxpayers *via* the Korea Basic Science Institute [IBS-R022-A2-2022].

## Author Contributions

KY: Conceptualization, Investigation, Analysis, Writing (original draft), Writing (review and editing)

BW: Conceptualization, Investigation, Analysis, Writing (review and editing)

NM: Investigation, Writing (review and editing)

AS: Investigation

AW: Conceptualization, Investigation, Writing (review and editing)

AG: Conceptualization, Analysis, Writing (review and editing), Funding acquisition

AV: Conceptualization, Analysis, Writing (original draft), Writing (review and editing), Funding acquisition

