## Supplemental Text and Figures for "Disparate roles for *C. elegans* DNA translocase paralogs RAD-54.L and RAD-54.B in meiotic prophase germ cells"

This PDF file includes Supplemental Text, SI references, Table S1, and Figures S1-S10.

### <u>Supplementary Information Text</u>

#### Supplemental Materials and Methods

##### *C. elegans* Genetics

The following strains were used:

| | |
|---|---|
| AV695 | *meIs8[pie-1p::GFP::cosa-1::unc-119(+)] II; mnT12 (X; IV)* |
| AV863 | *nbs-1(me106)/mnC1 II* |
| AV1158 | *him-6(jf93[him-6::HA]) rad-54.B(gt3402[rad-54.B::GFP]) IV* |
| AV1179 | *rad-54.L(me98)/tmC18[dpy-5(tmIs1236)] I* |
| AV1189 | *spo-11(me44) rad-54.B(gk340656)/tmC5[F36H1.3(tmIs1220)] rad-54.B(gk340656) IV* |
| AV1190 | *rad-54.L(me98)/tmC18[dpy-5(tmIs1236)] I; spo-11(me44) rad-54.B(vc205209)/tmC5[F36H1.3(tmIs1220)] rad-54.B(gk340656) IV* |
| AV1191 | *rad-54.L(me98) I; opIs257 [rad-54.Lp::rad-54.L::YFP::rad-54.L 3'UTR + unc-119(+)]* |
| AV1196 | *rad-54.L(me98)/tmC18[dpy-5(tmIs1236)] I; rad-54.B(gk340656) IV* |
| AV1199 | *spo-11(me44)/tmC5[F36H1.3(tmIs1220)] IV* |
| AV1203 | *rad-54.L(me98)/tmC18[dpy-5(tmIs1236)] I; spo-11(me44)/ tmC5[F36H1.3(tmIs1220)] IV* |
| AV1214 | *rad-54.L(me98) I; opIs257 [rad-54.Lp::rad-54.L::YFP::rad-54.L 3'UTR + unc-119(+)]; rad-54.B (gk340656) IV* |
| AV1215 | *rad-54.B(gk340656) IV; egl-1(n1084) chk-2(gk212)/egl-1(n1084) yIs34 oxTi633 V* |
| AV1216 | *rad-54.B(gk340656) IV; egl-1(n1084) yIs34 oxTi633 V* |
| AV1238 | *rad-54.L(me139)/tmC18[dpy-5(tmIs1236)] I; ieSi21[sun-1::mRuby] IV* |
| AV1239 | *rad-54.L(me139)/tmC18[dpy-5(tmIs1236)] I; ieSi11[EmGFP::syp-3] II; rad-54.B(gk340656) IV* |
| AV1284 | *rad-54.L(me177)/tmC18[dpy-5(tmIs1236)] I* |
| AV1292 | *rad-54.L(me177)/tmC18[dpy-5(tmIs1236)] I; spo-11(me44) rad-54.B(gk340656)/tmC5[F36H1.3(tmIs1220)] rad-54.B(gk340656) IV* |
| CB4856 | *C. elegans* wild isolate 'Hawaiian' |
| CB5584 | *mIs12[myo-2p::GFP + pes-10p::GFP + F22B7.9p::GFP] II* |
| TG3319 | *rad-54.B(gk340656)* |
| TG4580 | *rad-54.B(gt3328) IV* |
| TG4080 | *rad-54.B(gk340656) IV; meIs8[pie-1p::GFP::cosa-1::unc-119(+)] II* |
| TG4252 | *rad-54.B(gk340656) IV; meIs8[pie-1p::GFP::cosa-1::unc-119(+)] II; mnT12 (X; IV)* |
| XSW933 | *rad-54.B(gk340656) IV;* 'Hawaiian' V . (This strain was derived from inbreeding following an intercross between TG3319 and CB4856 to yield a strain carrying the Hawaiian-derived alleles of chromosome V SNP markers.) |

**CRISPR-Cas9 Gene Editing**

*rad-54.B(gt3402[rad-54.B::GFP])* was generated by CRISPR-Cas9 gene editing using sgRNA(GAATGCCGGGTTGTCGCAGG). The PU6::sgRNA template was generated by PCR as described in (Ward, 2015). 5' and 3' homology arms were generated from N2 genomic DNA by PCR (primer sequences below) and then inserted into the GFP^SEC^3xFlag vector pDD282 using Gibson assembly (New England BioLabs). A mixture of pDD162 (Peft-3::Cas9 , 50 ng/μL), pCFJ90 ( Pmyo-2 :: mCherry , 2.5 ng/μL) and pCFJ104 (Pmyo-3::mCherry , 5 ng/μL) plasmids (Dickinson et al., 2013), together with pU6-rad-54B sgRNA (50 ng/μL) and rad-54B:gfp repair templates (50 ng/μL), was microinjected in N2 young adult worms, followed by the screening, SEC excision and DNA sequencing as previous described in (Dickinson et al., 2015).

PCR primers for generation of 5' homology arm:

F: acgttgtaaaacgacggccagtcgccggcaCATCGATGCTCCTGAGGCTCCCGATGCTCC

R: CATCGATGCTCCTGAGGCTCCCGATGCTCCGgagcatcgggagcctcaggagcatcgatg

PCR primers for generation of 3' homology arm:

F: CGTGATTACAAGGATGACGATGACAAGAGATGATATTGATCTGTTGAATTTGTTT

R: ggaaacagctatgaccatgttatcgatttcCGCCGTAAATCTACCCTC

*rad-54.L(me177)* was generated using CRISPR-Cas9 gene editing as in (Paix et al., 2015). An alignment of *S. cerevisiae* Rad54 and *C. elegans* RAD-54.L was used to design the missense mutation in *rad-54.L(me177)*, which encodes RAD-54.L(K238R) with the identical amino acid substitution at the corresponding position present in the *S. cerevisiae* ATPase-dead mutant Rad54(K341R) (Clever et al., 1999; Petukhova et al.,

1999). sgRNA (ATGGCAGATGAAATGGGTCT) and ssDNA repair template (ATTAATATTCCCGAATTTCATGGATGTATTATGGCAGATGAAATGGGTCTTGGAAG GACACTTCAATGCATTTCTCTACTTTGGACACTTCT) (IDT) were used for microinjection into the germlines of WT animals. Animals carrying the edit were identified using PCR (F: GCGTCCGCATCAAAGAGATG, R: TCGAAACTGTTGGAC ACGCA) followed by a CviKI1 restriction enzyme digest. The edit was confirmed by Sanger sequencing.

*rad-54.B(gt3328)* was generated by CRISPR-Cas9 gene editing as described previously using Cas9 protein (0.25ug/ul), tracrRNA (0.02ug/ul), crRNA (0.02ug/ul) (5'-AACCACGTCAGCAGTCACTG[TGG]-3'), *rol-6(su1006)* (40ng/ul) (Dokshin et al., 2018) and a ssODN (0.11ug/ul) was used as a repair template with 40 bp homology arms flanking a 43bp universal STOP-in cassette (Wang et al., 2018) (5'GAACAGGATATTC TCGATCGAAAAACCACGTCAGCAGTCAGGGAAGTTTGTCCAGAGCAGAGGTGACT AAGTGATAAGCTAGCCTGTGGACACGAGATTCGCGAGACATCTCCGACCTCATCA G-3'). The final concentration of the components of the injection mix are indicated. F1 rollers were screened by PCR using the following primers (5'-CGTCTGTAAACTCTTT CCGAG-3' and 5'-GGGTGGAGCCTAAATTTAAG-3'). Successful insertion resulted in a 513 bp fragment (WT fragment 470 bp) and was verified by sequencing.

**Immunofluorescence**

Immunofluorescence experiments using whole-mount gonads or spread nuclei were conducted as in (Martinez-Perez and Villeneuve, 2005; Pattabiraman et al., 2017;

Woglar and Villeneuve, 2018); worms were dissected at 22-30 hours post L4. The following primary antibodies were used: chicken anti-HTP-3 (1:1000, (MacQueen et al., 2005)), chicken anti-GFP (1:250, (Abcam)), mouse anti-GFP (1:500, (Roche)), rabbit anti-GFP (1:500, (Yokoo et al., 2012)), mouse anti-HA (1:1000, (Clone 16B12; Covance)), rabbit anti-MSH-5 (1:10000, (SDIX)), rabbit anti-RAD-51 (1:500, (Colaiácovo et al., 2003)), rat anti-RAD-51 (1:200, (Rosu et al., 2013)), guinea pig anti-SUN-1 pS24 (1:700, (Penkner et al., 2009)), guinea pig anti-SYP-1 (1:200, (MacQueen et al., 2002)). Secondary antibodies used were Alexa Fluor 405 (1:100), 488 (1:400), 555 (1:400), and 647 (1:200)-conjugated goat antibodies raised against the appropriate species (Life Technologies).

**Imaging**

Image acquisition and processing of whole-mount gonads and spread nuclei were conducted as in (Roelens et al., 2019; Woglar and Villeneuve, 2018). All images were acquired on a DeltaVision OMX Blaze microscope with a 100x NA 1.4 NA objective, with 200 nm spaced z-stacks for widefield images and 125 nm spaced z-stacks for SIM images. For widefield images, images were deconvolved and registration corrected using SoftWoRx software. Individual fields of view were assembled together for each gonad using the "Grid/Collection stitching" FIJI plugin (Preibisch et al., 2009). For display, a single layer of nuclei was maximum-projected using FIJI, except for **Fig 5A**, **6A**, and **S8**, in which a single z-slice at the center of the nuclear layer is displayed. For SIM images, images were reconstructed, registration corrected, and maximum projected using SoftWoRx software. Brightness and contrast were adjusted in FIJI for display of both widefield and SIM images.

**Focus count quantification pipeline**

Z-stack images of whole gonads were cropped to include only a single layer of germ cell nuclei. These cropped images ("cropped Z-stacks") were max projected, and nuclei that were well-separated from each other were manually segmented using HTP-3 signal and saved as polygon-shaped ROIs in FIJI software. These ROIs were then overlayed on the cropped Z-stacks, and the area outside of the ROIs was cleared of all signals. Then, the 3D Maxima Finder plugin on ImageJ was used to identify immunofluorescence foci and determine their peak brightness and the 3D positions of their maxima (Ollion et al., 2013). For 3D Maxima Finder parameters, the "Minimum Peak Height" was determined empirically, starting at the max value of background fluorescence as determined by manual sampling, and iteratively running the plugin with different "Minimum Peak Height" values to minimize the numbers of false positive and false negative foci identified. All other parameters ("Radius xy", "Radius z", "Noise") were kept at the default settings, 1.5, 1.5, and 100, respectively. From the 3D Maxima Finder output table of foci with their xyz peak positions and peak heights, foci were computationally assigned to ROIs corresponding to individual nuclei based on their positions. Nucleus-focus assignments were manually checked to remove overlap to generate the final foci counts. The xy position of each nucleus ROI was used to approximate its position within the region of quantification, *i.e.* the x-axis values in **Fig 1E and 2B**.

For whole-gonad quantifications (**Fig 1E**), this analysis was performed across the entire gonad. For quantification in early pachytene nuclear spreads (**Fig 2B, 2E, 5E, S9**), the early pachytene zone was defined as starting at the cell row in which most nuclei have a RAD-51 focus and ending at the cell row in which most nuclei have 6 MSH-5 or HIM-6 foci. For quantification in late pachytene nuclear spreads (**Fig 2E**), the late pachytene zone was defined as starting at the cell row in which most nuclei have 6 MSH-5 foci and ending before the diplotene stage when HTP-3 axis staining loses its linear appearance. For both cases, the appropriate region was cropped and used for the quantification pipeline above. For quantification of RAD-51 foci in *rad-54.L* and *rad-54.B* mutants (**Fig 3B-E, S6**), a 10 cell-row wide zone corresponding to the region of peak accumulation of RAD-51 foci was manually identified and cropped before performing focus count quantification as described above.

**Meiotic crossover (CO) distribution assay using SNP mapping strategy**

Meiotic CO distribution was assayed as described (Agostinho et al., 2013). TG3319(*rad-54.B(gk340656)*) was crossed to CB4856 to generate a new strain XSW933 homozygous for the *rad-54B(gk340656)* mutation as well as the CB4856-derived alleles at SNP sites, V-17.5, V-5, V5.8, V17.8, V25 (Davis et al., 2005) (**Table S1**). We then crossed TG3319 males to XSW933 young adult hermaphrodites, and crossed N2 males to CB4856 young adult hermaphrodites as a control. Following successful mating (~12 hours), L4 hermaphrodite worms of the $F_1$ generation from both crosses were individually picked and allowed to grow to the young adult stage. These hermaphrodites were mated to CB5584 males (carrying a pharyngeal GFP marker). After ~12 hours of mating, the $F_1$ animals were plated to lay eggs for two days. F1 animals were picked for single-worm lysis and genotyping by PCR. From the $F_1$ plates that were confirmed to be heterozygous for the CB4856-derived alleles of the chromosome *V* SNP markers , 256 $F_2$ cross-progeny L4 hermaphrodites (expressing the pharyngeal GFP marker) were picked. Single worm lysis was carried out and the V-17.5, V-5, V5.8, V17.8, V25 SNP sites were tested for presence of the CB4856-derived alleles by PCR. The primers used for the PCR are indicated in **Table S1**. The PCR reaction was set up at 94°C for 40 seconds, followed by 35 cycles of 40 seconds at 60°C and 1 minute at 72°C. This was followed by 10 minutes at 72°C. PCR products were digested with Takara Quickcut™ Dra I (Aha III) restriction enzyme according to the manufacturer's instruction, then resolved on a 2% agarose gel (Sangon Biotech) for 40 minutes.

**<u>Table S1</u>: Dra I SNP locations and primers sequences for CO distribution assay.**

| Location | primers |
|---|---|
| V -17.5 F | GCGACTGTCACAATCAAGA |
| V -17.5 R | CTGCTTGGCTTCCTCTA |
| V -5 F | GAGATTCTAGAGAAATGGACACCC |
| V -5 R | AAAAATCGACTACACCACTTTTAGC |
| V 5.8 F | CCTTATCTAGTAATTTGCCTGTTGT |
| V 5.8 R | ACATAAGCGCCATAACAAGTCG |
| V 17.8 F | GAAATTCAAATTTTTGAGAAACCC |
| V 17.8 R | TTCAGACCATTTTTAGAATATTCAGG |
| V 25.1 F | ACTTGACTCCTCTTTTCCATG |
| V 25.1 R | CTGCTAGCTCAAATACTCCC |

S1



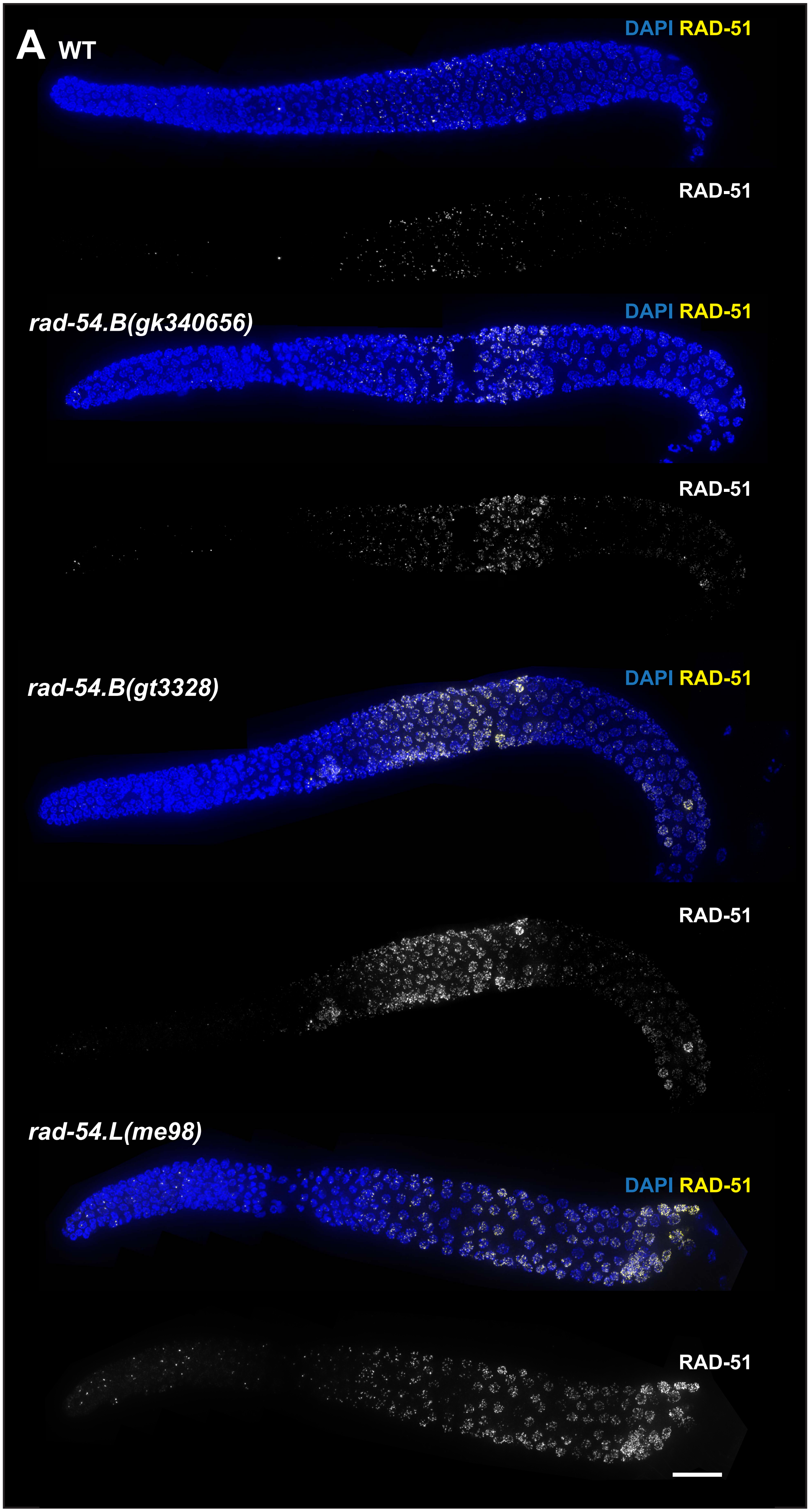


S1. **Hyperaccumulation of RAD-51 in *rad-54.B* mutants differs from the *rad-54.L* mutant.** A) Max-projected images of whole-mount gonads from indicated genotypes immunostained for RAD-51. While both *rad-54.B* and *rad-54.L* mutants have elevated levels of RAD-51 foci, in both *rad-54.B* mutants, elevated RAD-51 foci decline midway through meiotic progression, whereas RAD-51 foci continue to accumulate in the *rad-54.L* mutant. Scale bar represents 20 μm.

# S2

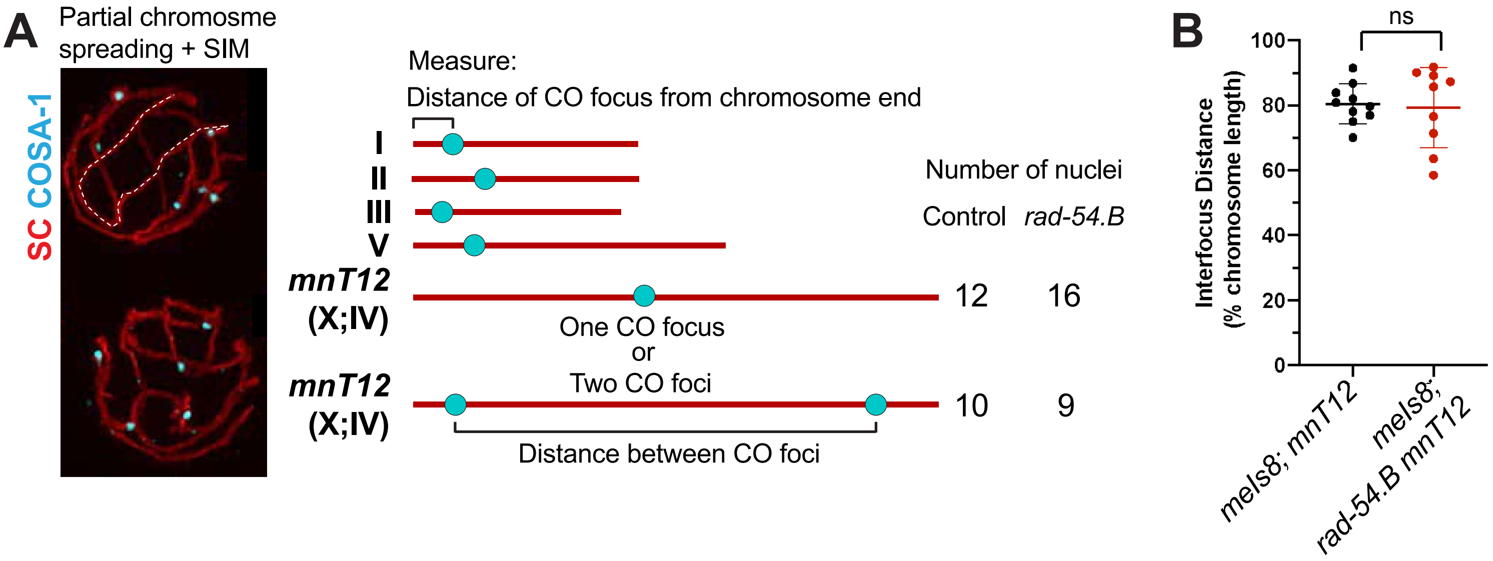


S2. **Normal spacing between CO-site foci in the *rad-54.B* mutant background.** A) Left: Examples of late pachytene nuclear spreads, imaged with SIM, that were used for chromosome tracing and quantification of positions of CO-designated recombination sites marked by immunolocalization of GFP::COSA-1 (cyan) and MSH-5 (not depicted). Middle: Schematic of the karyotype of the strains analyzed (control: *meIs8[pie-1p::GFP::cosa-1::unc-119(+)]; mnT12*, and *rad-54.B*: *meIs8; rad-54.B(gk340656) mnT12*) and the features quantified. Right: Numbers of nuclei assayed in control (*meIs8; mnT12*) and *rad-54.B* (*meIs8; rad-54.B mnT12*) backgrounds that had one or two CO foci on the *mnT12* fusion chromosome bivalent. B) Quantification of inter-focus distances on *mnT12* fusion chromosome bivalents that had 2 CO site foci, represented as a percentage of the total *mnT12* chromosome length. Control, n=10 chromosomes; *rad-54.B*, n=9 chromosomes. Statistical significance was assessed with a Mann Whitney test; ns, p>0.05.

# S3

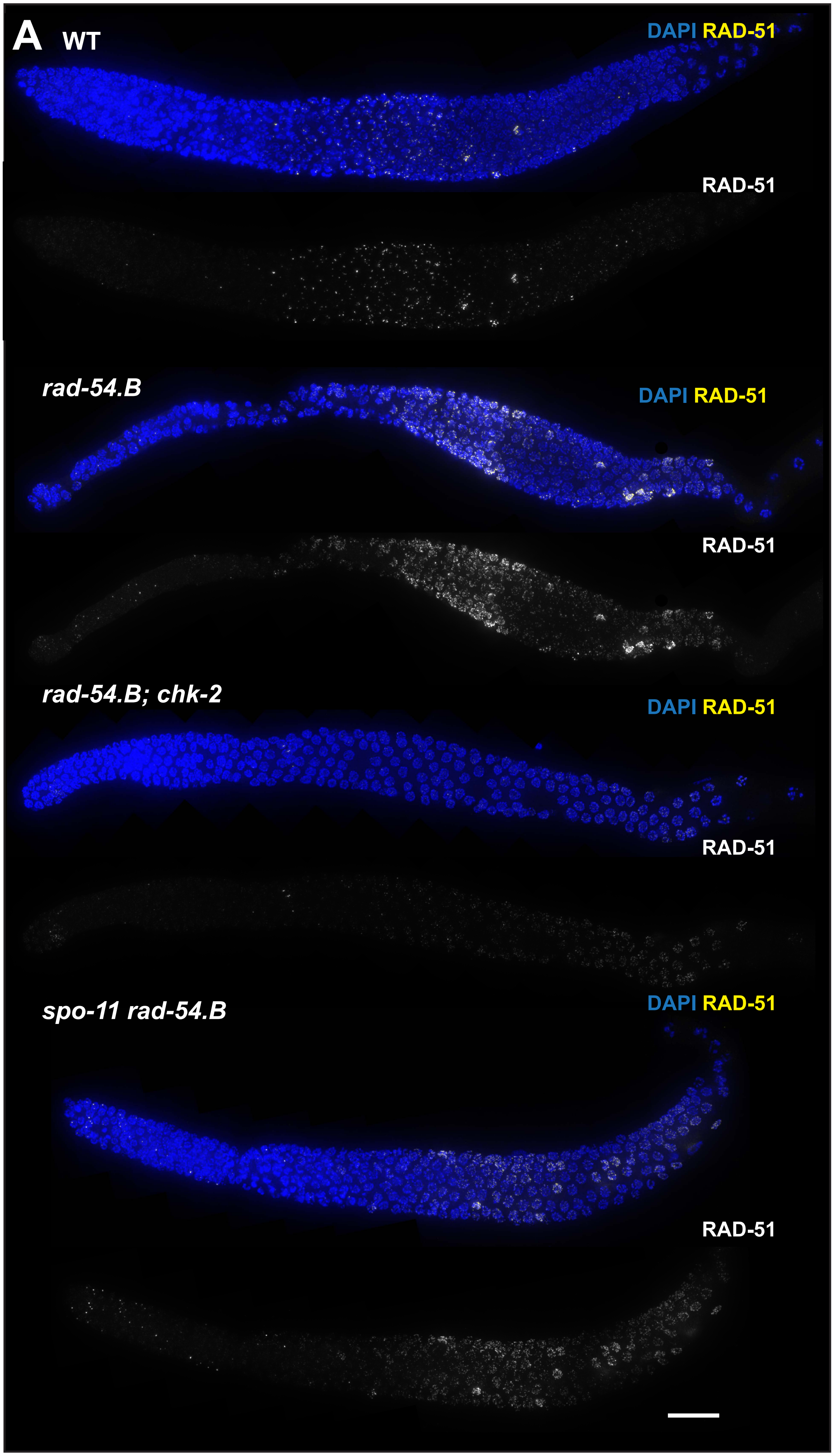


S3. **Hyperaccumulation of RAD-51 foci in *rad-54.B* background is dependent on CHK-2.** A) Max-projected images of whole-mount gonads from indicated genotypes immunostained for RAD-51. Full genotypes: WT, *rad-54.B(340656); egl-1 yIs34 oxTi633*, *rad-54.B; egl-1 chk-2*, *spo-11 rad-54.B*. We note that while there are some faint RAD-51 foci detected in the *rad-54.B; chk-2* mutant, these are far weaker, fewer in number and occur much later in prophase than the RAD-51 foci seen in WT, *rad-54.B*, or *spo-11 rad-54.B* germ lines, supporting the conclusion that the RAD-51 hyperaccumulation phenotype observed in rad-54.B mutants is dependent on CHK-2. Scale bar represents 20 µm.

# S4



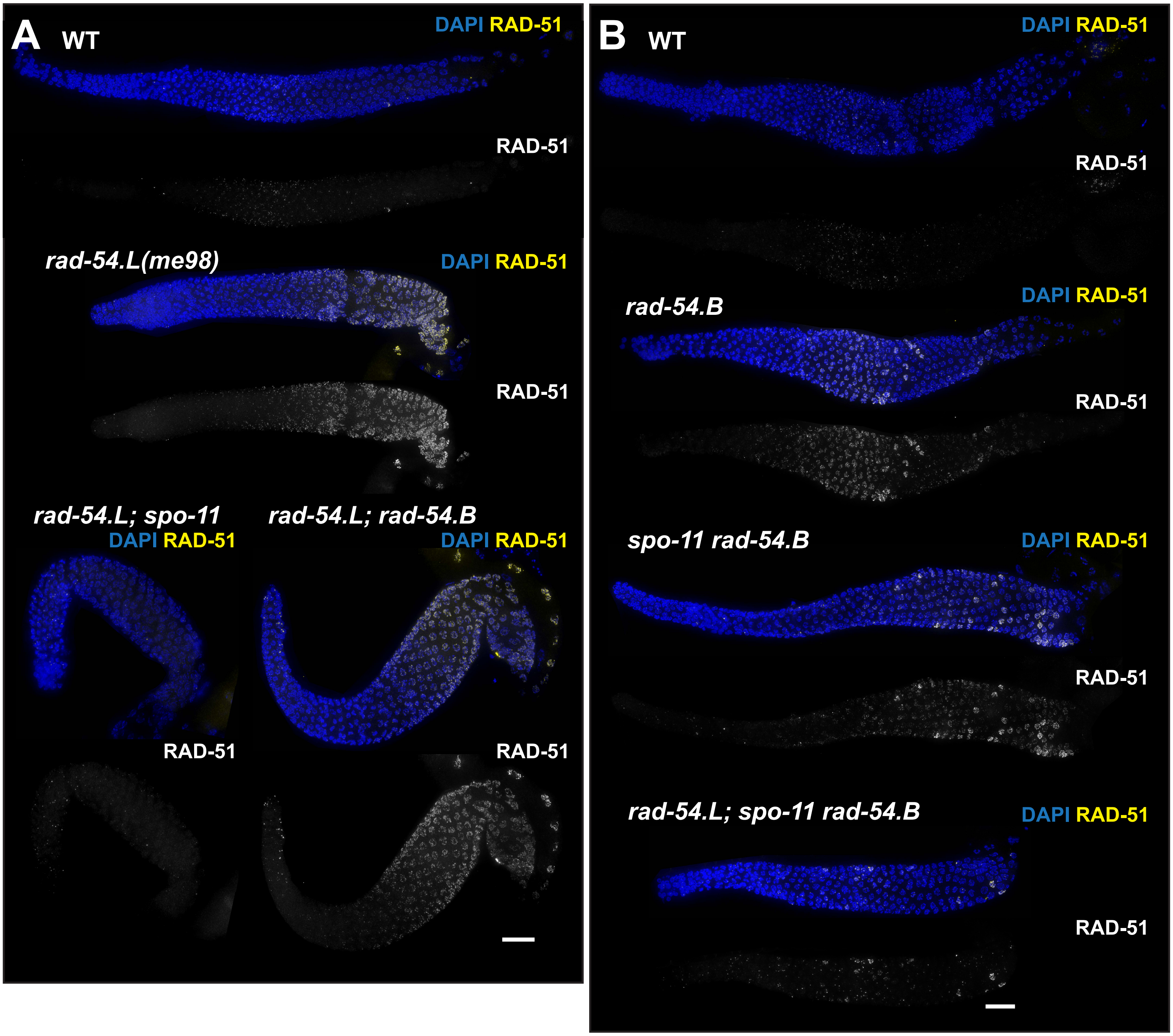


S4. **RAD-51 hyperaccumulates at unbroken DNA in the *rad-54.B* mutant.** A) and B) Max-projected images of whole-mount gonads from indicated genotypes immunostained for RAD-51. Scale bar represents 20 µm.

# S5



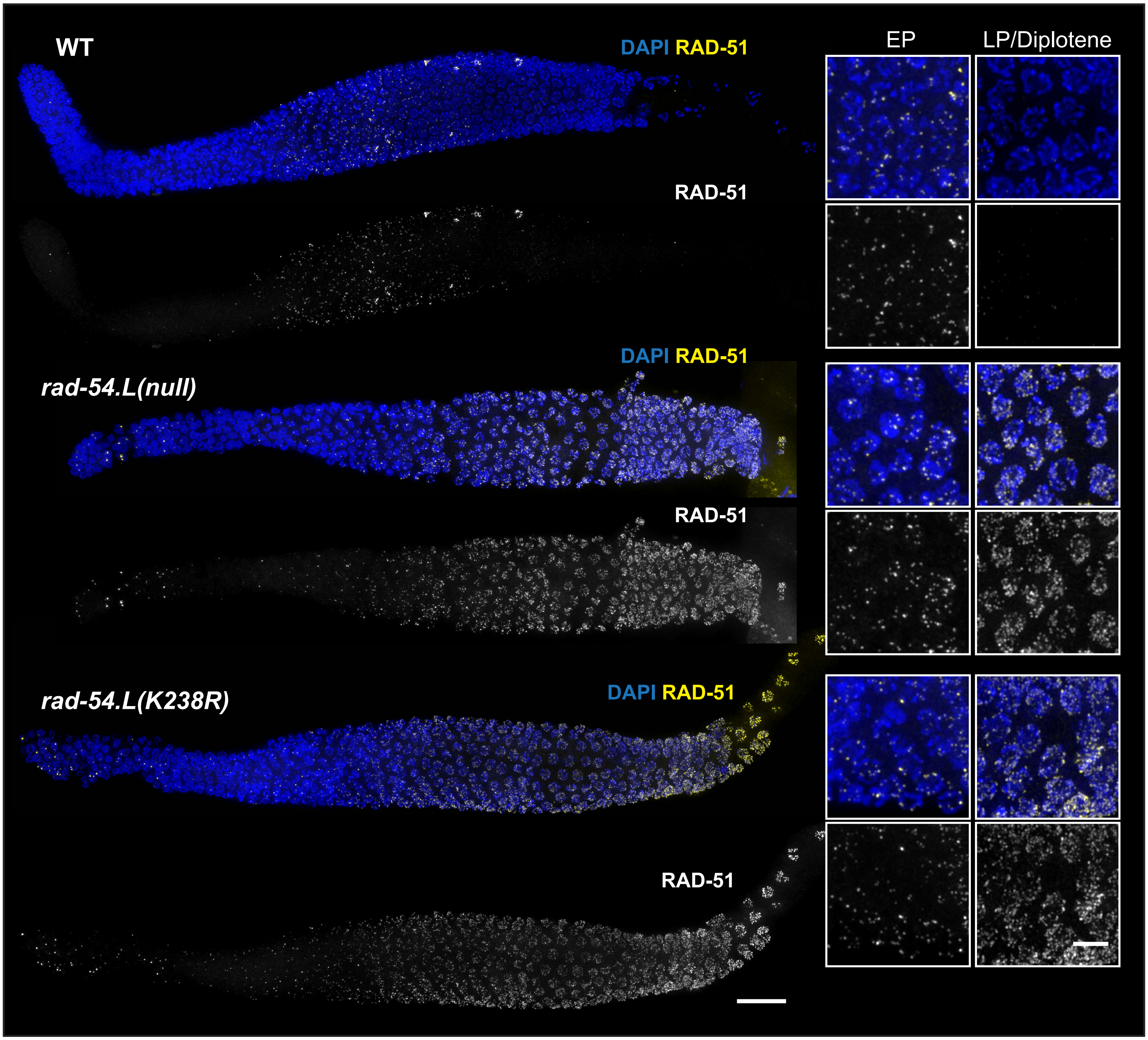


S5. **RAD-51 hyperaccumulates in *rad-54.L(K238R)* germ lines similarly to *rad-54.L(me98)(null)* germ lines.** Max-projected images of whole-mount gonads from indicated genotypes immunostained for RAD-51. Scale bar represents 20 µm. Insets at right depict zoomed-in fields of nuclei from early pachytene (EP) and late pachytene (LP)/diplotene stages. Scale bar in insets, 5 µm.

# S6



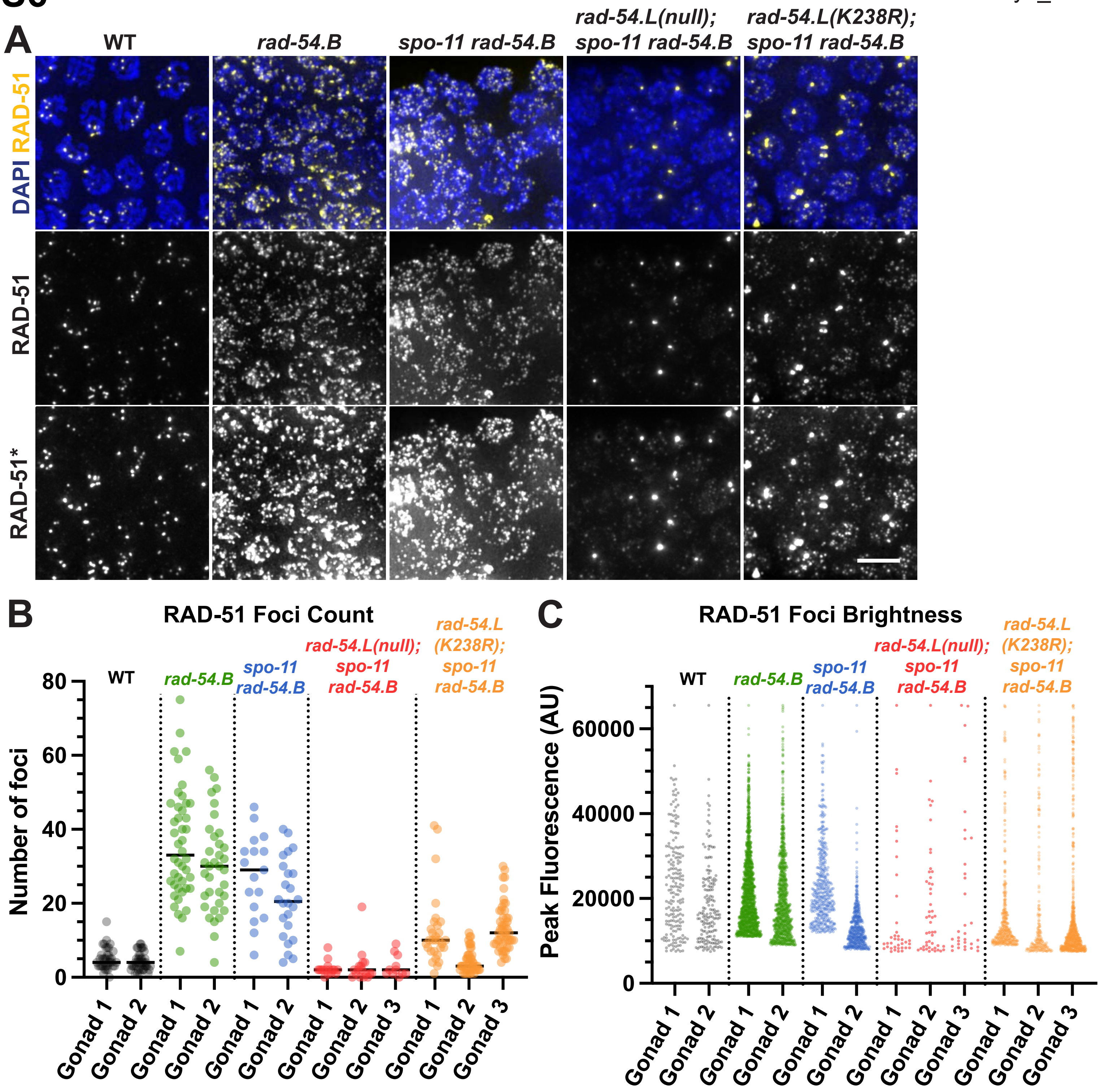


Figure legend on following page.

S6**. Effect of *rad-54.L(K238R)* ATPase-dead mutant on SPO-11-independent accumulation of RAD-51.** A) Representative fields of nuclei from max-projected images of whole-mount gonads, taken from the zones of maximum accumulation of RAD-51 foci for the indicated genotypes; scale bar represents 5 μm. Comparison of images reveals: 1) a substantial attenuation of SPO-11-independent RAD-51 hyperaccumulation in *rad-54.L(K238R); spo-11 rad-54.B* relative to *spo-11 rad-54.B*, and 2) more abundant residual SPO-11-independent RAD-51 foci in *rad-54.L(K238R); spo-11 rad-54.B* relative to *rad-54.L(null); spo-11 rad-54.B*. Bottom row depicts overexposure of RAD-51 signal (RAD-51*), which reveals very faint nuclear foci in *rad-54.L(null); spo-11 rad-54.B* germ cells that are dimmer than those detected in *rad-54.L(K238R); spo-11 rad-54.B* and are below the threshold intensity required for reliable quantification. B) Quantification of RAD-51 foci for the genotypes shown in panel A. RAD-51 foci counts were conducted in the zone of maximum accumulation of RAD-51 foci for each genotype. Each circle represents a nucleus; lines indicate median values. Numbers of <u>nuclei</u> analyzed (n) and median numbers of foci (m) were as follows: WT gonad 1, n =37, m=4, gonad 2, n=40, m=4; *rad-54.B* gonad 1, n=44, m=33, gonad 2, n=35, m=30; *spo-11 rad-54.B* gonad 1, n=17, m=29, gonad 2, n=24, m=20.5; *rad-54.L(null); spo-11 rad-54.B* gonad 1, n=16, m=2, gonad 2, n=17, m=2, gonad 3, n=11, m=2; *rad-54.L(K238R); spo-11 rad-54.B* gonad 1, n=30, m=10, gonad 2, n=45, m=3, gonad 3, n=50, m=12. C) Peak intensity of RAD-51 foci detected in the zone of maximum accumulation for each genotype. Each point in the scatter plot represents a RAD-51 focus. Note that quantification pipeline was run in some gonads with a higher threshold peak value to minimize false positive peaks (e.g. peaks identified in off-focus z-planes or in background control areas). Numbers of <u>foci</u> analyzed (n) were as follows: WT gonad 1, n=186, gonad 2, n=169; *rad-54.B* gonad 1, n=1587, gonad 2, n=1047; *spo-11 rad-54.B* gonad 1, n=467, gonad 2, n=509; *rad-54.L(null); spo-11 rad-54.B* gonad 1, n=37, gonad 2, n=49, gonad 3, n=33; *rad-54.L(K238R); spo-11 rad-54.B* gonad 1, n=370, gonad 2, n=201, gonad 3, n=669.

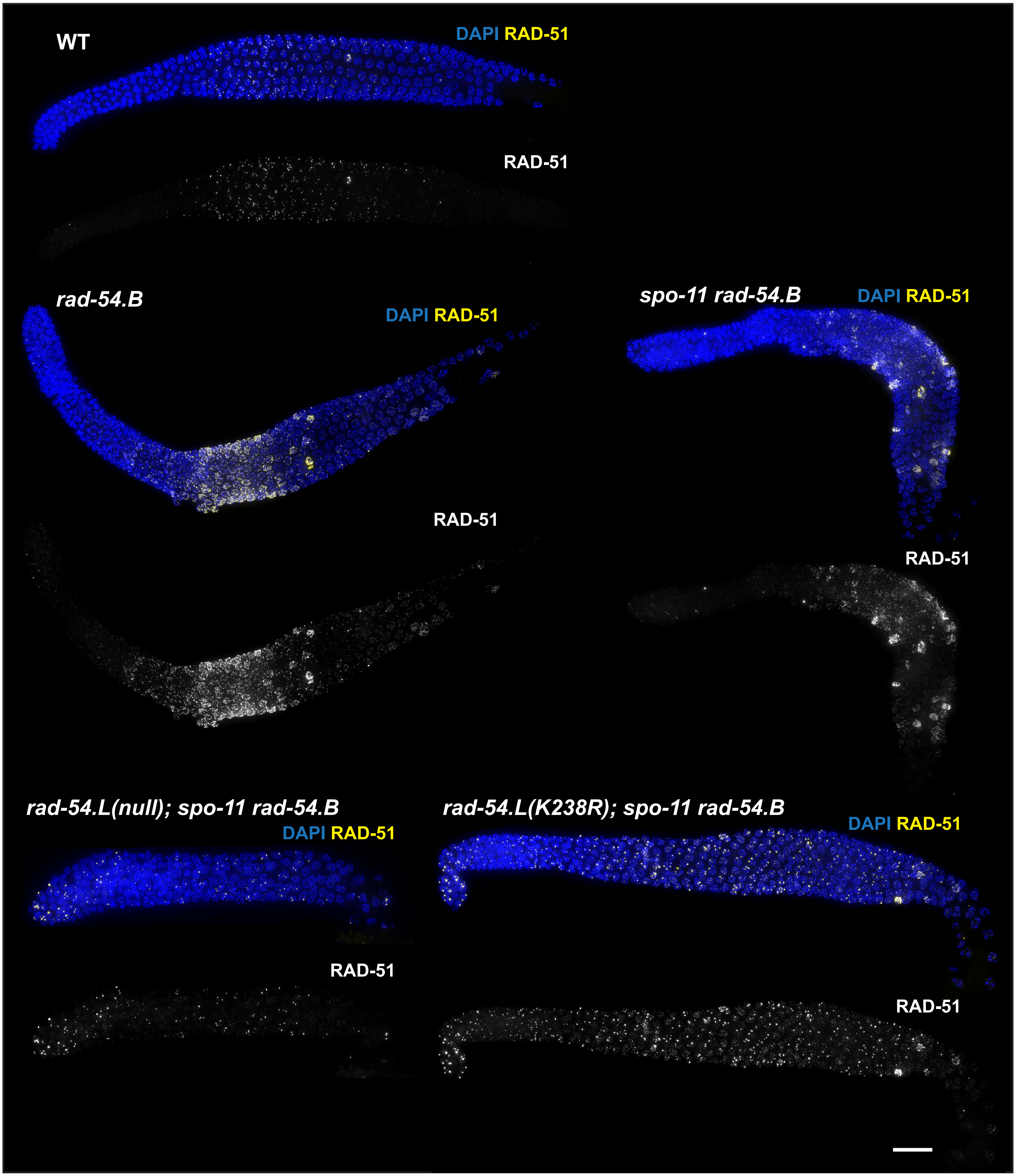


S7. **Effect of *rad-54.L(K238R)* ATPase-dead mutant on SPO-11-independent accumulation of RAD-51 (continued).** Max-projected images of whole-mount gonads from indicated genotypes immunostained for RAD-51. Scale bar represents 20 µm.

# S8

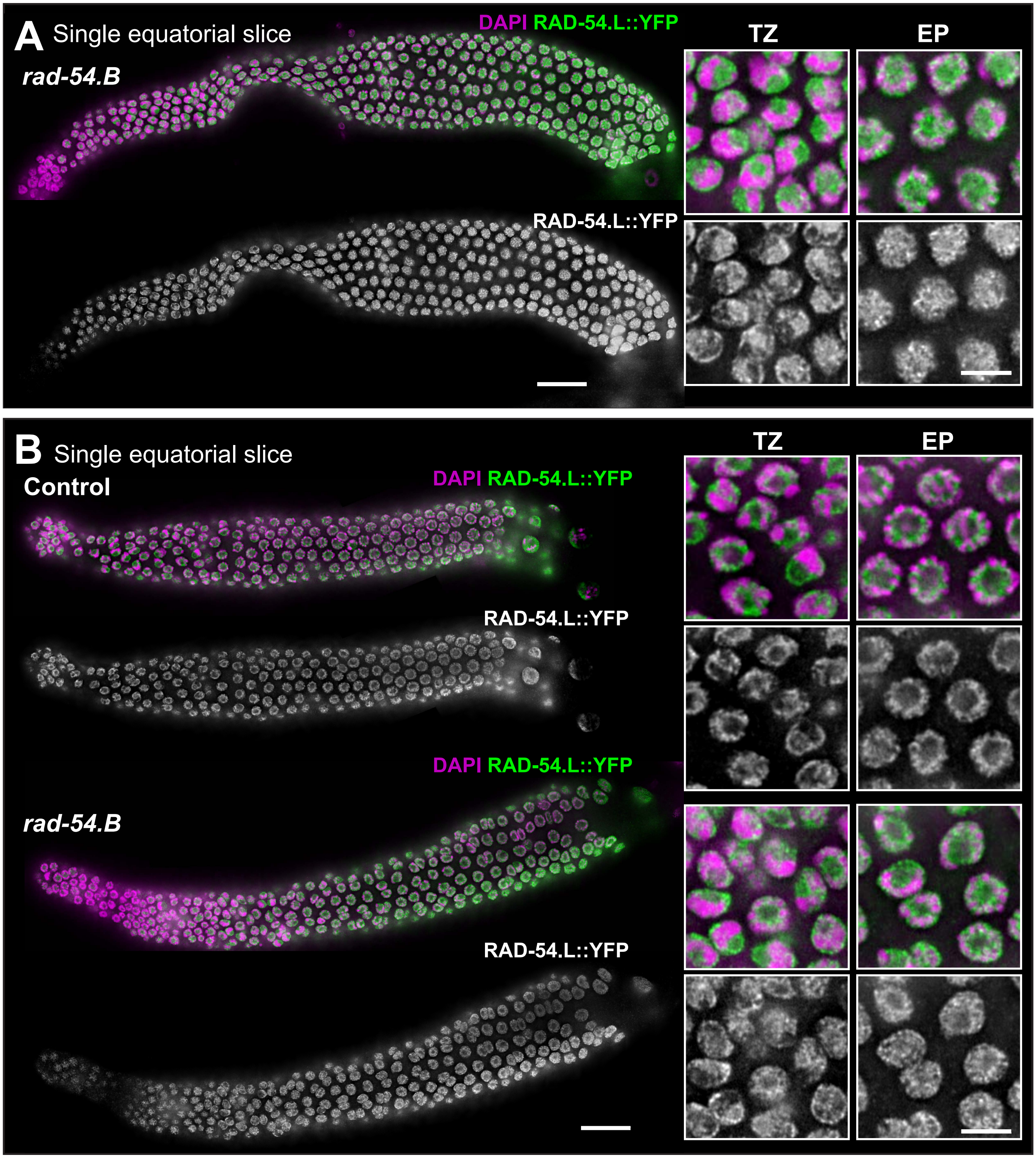


S8. **RAD-54.L::YFP localization.** A) Image of whole-mount gonad from a *rad-54.L(null); opIs257[rad-54.Lp::rad-54.L::YFP::rad-54.L 3'UTR + unc-119(+)]; rad-54.B* worm immunostained for RAD-54.L::YFP. The image represents a single z-slice showing an equatorial view of nuclei (instead of a max-projection). Scale bar represents 20 µm. Insets at right depict zoomed-in fields of nuclei from the transition zone (TZ) and early pachytene zone (EP). Scale bar in insets, 5 µm. B) Image of whole-mount gonad immunostained for RAD-54.L::YFP from worms of the indicated genotypes. Full genotypes are as detailed in Fig. 5A and S8A. The image represents a single z-slice showing an equatorial view of nuclei (instead of a max-projection). Unlike in Fig. 5A and S8A, these images show examples of gonads in which RAD-54.L::YFP signal is not observed in the nucleolus. Scale bar represents 20 µm. Insets at right depict zoomed-in fields of nuclei from the transition zone (TZ) and early pachytene zone (EP). Scale bar in insets, 5 µm.

# S9

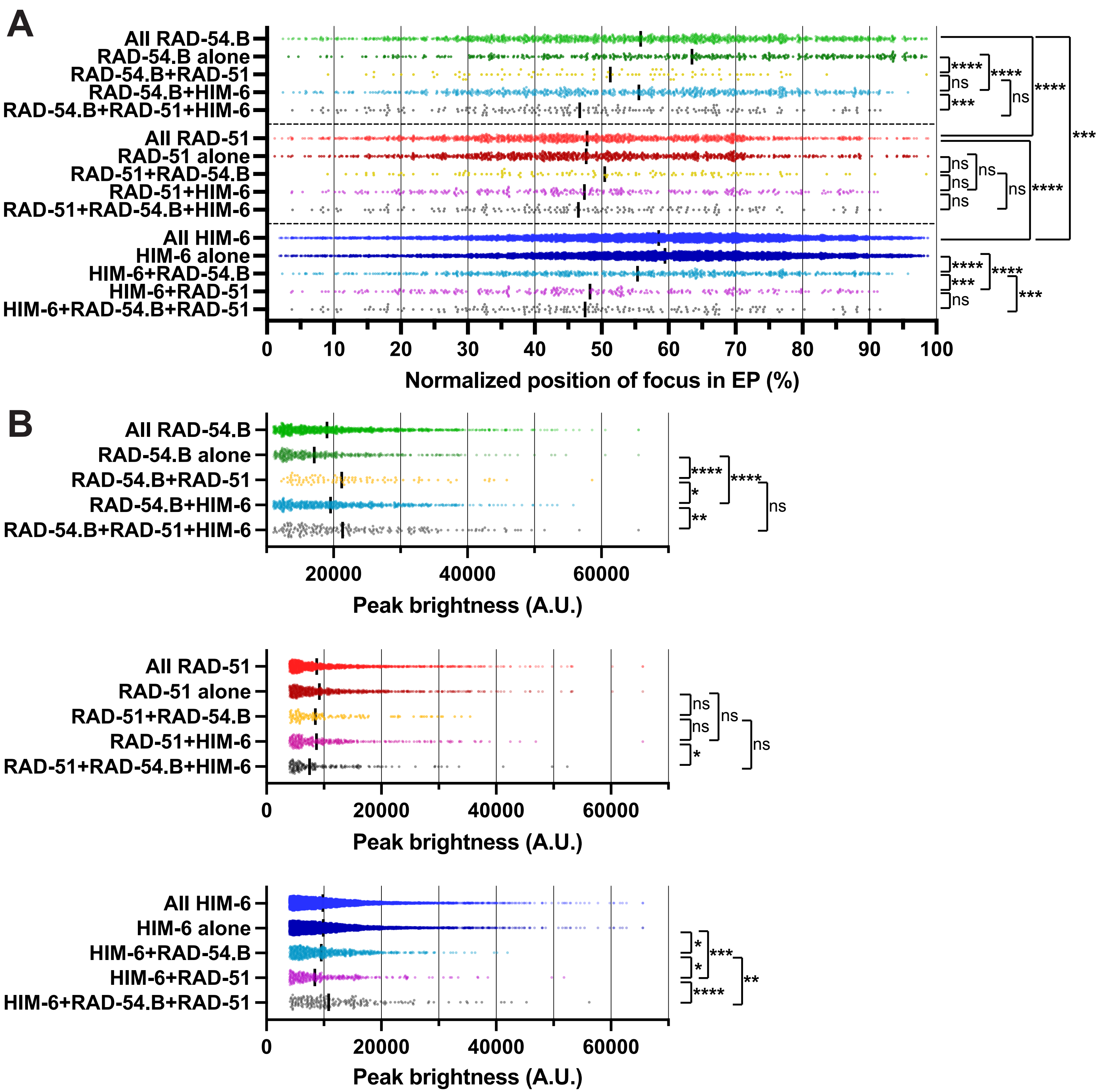


Figure legend on following page.

S9. **Positions and peak brightness of RAD-54.B::GFP, RAD-51, and HIM-6 foci in early pachytene nuclear spreads.** A) Positions of early pachytene RAD-54.B::GFP, RAD-51, and HIM-6 foci with the indicated colocalization pattern, depicted as a percentage of the length of the total early pachytene zone. Early pachytene zone was defined as starting at the cell row in which all nuclei have a RAD-51 or HIM-6 focus (0%) and ending immediately before the cell row in which all nuclei have bright, CO-site associated HIM-6 foci (100%). Each point in the scatter plot represents a focus with the indicated colocalization pattern (alone, or with one or both other DSBR proteins) and the line indicates the median value. Statistical significance was assessed with a Mann Whitney test; ns, $p>0.05$; ***, $p<0.001$; ****, $p<0.0001$. B) Peak brightness values of early pachytene RAD-54.B::GFP, RAD-51, and HIM-6 foci with the indicated colocalization pattern. Each point in the scatter plot represents a focus with the indicated colocalization pattern and the line indicates the median value. Statistical significance was assessed with a Mann Whitney test; ns, $p>0.05$; *, $p<0.05$; **, $p<0.01$; ***, $p<0.001$; ****, $p<0.0001$.

# S10



**Chromsome V domains**

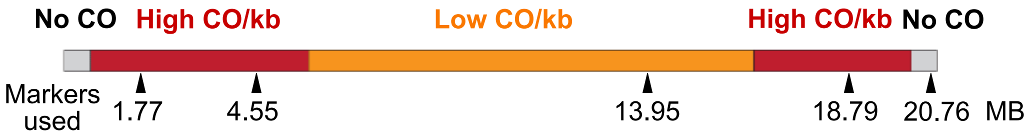


| | Genotype | Interval 1 | Interval 2 | Interval 3 | Interval 4 | NCO | P-value |
|---|---|---|---|---|---|---|---|
| Expt 1 | WT | 29 | 18 | 34 | 12 | 99 | 0.034 |
| | *rad-54.B* | 18 | 32 | 23 | 12 | 105 | |
| Expt 2 | WT | 30 | 23 | 55 | 21 | 127 | 0.003 |
| | *rad-54.B* | 14 | 40 | 34 | 15 | 153 | |

S10. **The *rad-54.B* mutant exhibits a modestly altered distribution of COs**. Top: schematic of chromosome V, with chromosome domains as defined by recombination assays (Rockman and Kruglyak, 2009) depicted by the colored boxes, and SNP markers used indicated with arrowheads below. Bottom: numbers of COs detected in each interval, and numbers of non-CO (NCO) products, from two independent experiments. P-value comparing CO distribution in WT vs *rad-54.B(gk340656)* was calculated using a chi-square test for independence.
